# A balance of metabolism and diffusion articulates a gibberellin hormone gradient in the *Arabidopsis* root

**DOI:** 10.1101/2025.03.24.643304

**Authors:** Kristian B. Kiradjiev, J. Griffiths, Alexander M. Jones, Leah R. Band

## Abstract

The plant hormone gibberellin (GA_4_) regulates numerous developmental processes. Within the root, GA_4_ controls growth, in part, by controlling the extent of cell elongation. The nlsGPS1 FRET biosensor revealed a GA_4_ gradient within the *Arabidopsis* root growth zones, with GA_4_ levels correlating with cell length. We developed a multiscale mathematical model to understand how biosynthesis, catabolism and transport create the GA_4_ distribution within the root growth zones. The model showed that phloem delivery of the biosynthetic intermediate GA_12_ contributes to higher levels of bioactive GA_4_ in the elongation zone, with the GA_4_ synthesis pattern being further modified by local GA_12_ synthesis in the quiescent centre region and the spatial distribution of biosynthesis enzymes (GA20ox and GA3ox). Model predictions revealed that whilst GA20ox and GA3ox transcript is present throughout the growth zones, these enzymes are inactive in the dividing cells, which explains steep GA_4_ gradients observed in GA20ox and GA3ox over-expression lines, and improves agreement between model predictions and data in wildtype, *ga20ox* and *ga3ox* lines. The model revealed that the GA_4_ gradient also depends a balance of diffusion through plasmodesmata and catabolism. Both model predictions and biosensor data demonstrated that plasmodesmatal diffusion enables a more gradual GA_4_ gradient, with higher diffusion antagonizing the GA_4_ gradient. Model predictions suggested that catabolism limits GA_4_ levels, which we validated via biosensor imaging in the *ga2oxhept* mutant. We concluded that GA_4_ distribution mediates root growth programming via local GA_4_ synthesis combining with diffusion and catabolism to create a spatial gradient that provides positional information and patterns cell elongation.

## Introduction

The plant hormone gibberellin (GA_4_) controls numerous developmental processes and environmental responses. In *Arabidopsis thaliana*, GA_4_ controls root growth by regulating both cell production in the meristem (the region of dividing cells close to the root apex) [37, 2], and cell growth in the adjacent elongation zone (EZ) (where cells undergo rapid elongation) [36]. It is well established that plants with reduced GA_4_ biosynthesis (due to mutations in key enzymes) have reduced root growth, due to both a shorter meristem [37, 2] and a shorter elongation zone [36]; however, we lack mechanistic understand of how these mutations affect the GA_4_ distribution and how the GA_4_ distribution mediates the growth responses.

The GA_4_ FRET biosensor, nlsGPS1, revealed a GA_4_ gradient within the Arabidopsis root growth zone, with GA_4_ levels being lowest in the division zone and increasing as cells begin rapid elongation [27]. To understand the mechanisms controlling this root GA_4_ gradient, we created an initial multicellular mathematical model of GA_4_ dynamics within the root growth zones [28]. The model predicted that the GA_4_ gradient is due to low biosynthesis in the meristem, and an increase in biosynthesis towards the shootward end of the meristem [28]. While this initial model provided insights into a necessary differential in biosynthesis, it remained unclear how is this relates to the distribution of GA_4_ metabolism enzymes and whether cell-to-cell transport also influences the GA_4_ gradient.

There are numerous cell-scale processes that may affect the GA_4_ distribution across the root growth zone. Bioactive GA_4_ is synthesised through a series of oxidation steps, mediated by GA20oxidases (GA20ox) and GA3oxidases (GA3ox), from the precurssor GA_12_. This GA_12_ is both delivered to the root via the phloem [26] and locally synthesised [28]. The enzymes involved in the downstream GA_4_ metabolism (i.e. the combination of biosynthesis and catabolism) exhibit spatial variations in expression level [23, 28] which may lead to GA_4_ gradients. In addition, hormone distributions are influenced by transmembrane transport. Recent studies revealed that gibberellin metabolites are actively transported across cell membranes via NPF and SWEET proteins [18, 35, 9]. Furthermore, hormones diffuse between adjacent cells through plasmodesmata, which has been shown to influence hormone-related phenotypes [25], though how plasmodesmatal diffusion contributes to GA_4_ distribution is currently unknown.

Whilst the nlsGPS1 FRET biosensor has shown how perturbing the GA_4_ biosynthesis enzymes affects the GA_4_ gradient in the root growth zones [28], modelling is essential to infer how the GA_4_ gradient arises from dynamic processes at the cellular scale. By developing a multiscale mathematical model, we now reveal how metabolism and transport combine to articulate the GA_4_ gradient. We reveal how the GA_4_ gradient emerges from the combination of (i) GA_12_ delivered to the EZ via the phloem together with locally synthesised GA_12_, (ii) spatial patterns of GA20ox and GA3ox biosynthesis enzyme activity, (iii) plasmodesmatal diffusion and (iv) catabolism. The model predictions are refined and validated using FRET biosensor data in wildtype and lines with perturbed metabolism enzymes. Thus, our study reveals that GA_4_ distributes analogously to a classic ‘morphogen’ reminiscent of Wolpert’s French Flag pattern [41], with localized synthesis combining with diffusion and catabolism to create a spatial gradient that underlies growth regulation.

## Results

### Model description

To investigate the GA_4_ distribution in the *Arabidopsis* root growth zone, we created a cell-based mathematical model that predicts the GA_4_ concentration in a cell file. One end of the file is situated at the quiescent centre (QC), and the other is in the mature zone (MZ) ~1300 *µ*m from the QC. Thus, the file comprises the meristem, where cells elongate slowly and divide, the elongation zone (EZ), where cells rapidly elongate, and a portion of the mature zone (MZ), where cells have ceased elongation (Fig 1A). We modelled mature roots in which the sizes of the meristem and EZ are stable. We prescribed cell elongation from previous experimental measurements of the relative elongation rates (RER) [38] (SI Appendix, Fig. S1 and Table S1). We assumed that meristematic cells divide once they have approximately doubled in size; prescribing initial cell lengths based on measurements in [15]. When a cell reaches the end of the meristem (due to growth of the more rootward cells), it ceases division and begins rapid elongation.

**Figure 1:**
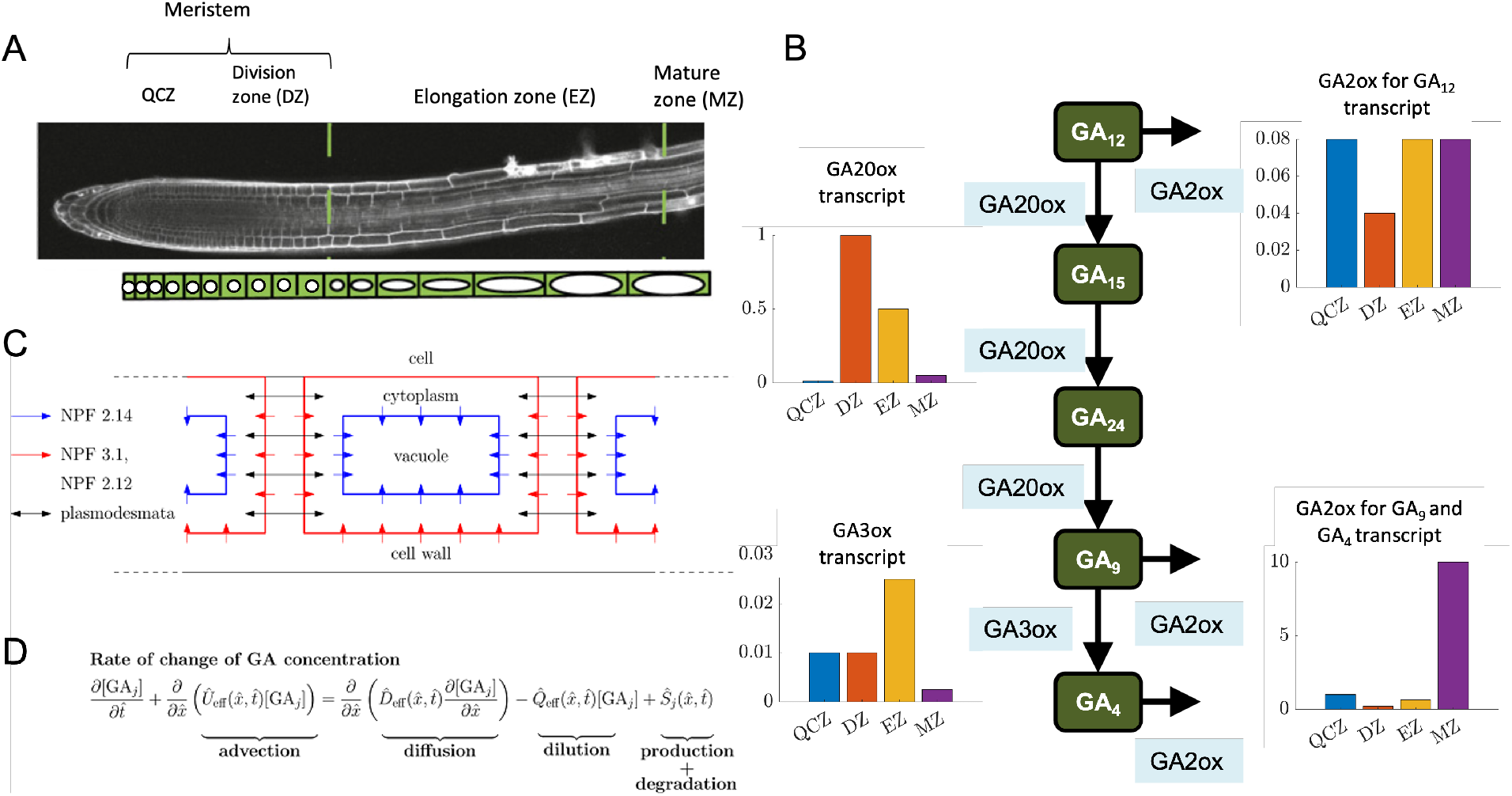
(**A**) Schematic of the cell-based model, representing the Arabidopsis root growth zones. (**B**) Schematic of the pathway downstream of GA_12_ that mediates the biosynthesis and catabolism of GA_4_, together with data showing the relative transcript levels of the enzymes, GA20ox, GA3ox, and GA2ox (showing separately the levels for GA2ox mediating GA_12_ degradation and for GA2ox mediating GA_9_ and GA_4_ degradation) [23, 28]. (**C**) Schematic focussing on a single cell showing the cell compartments, the transport components and the localisation of the NPF transporters. (**D**) Continuum governing equations for each metabolite, *j* = 12, 15, 24, 9, and 4. The equations for each metabolite are coupled through their production and degradation terms, 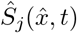, which represent the metabolic pathway depicted in Fig. 1B, and correspond to formulae given in SI Appendix section 5.

We simulated the GA_4_ concentration in each cell cytoplasm, cell vacuole and intercellular apoplastic compartment (SI Appendix, Fig. S2) (noting that the vacuole has recently been shown to contribute to GA_4_ distribution [9]). Within the meristem, the cells’ vacuolar volume fraction is constant (approximately 35% [11]), requiring both the cytoplasm and vacuole to expand with a relative growth rate equal to the RER of the cell. In the EZ, it is predominantly the vacuole that expands [11, 12]. Previous root models assumed that the cell elongation in the EZ was entirely due to vacuolar expansion [4, 28]; however, recent data suggest some cytoplasmic expansion in the EZ [11, 12]. Our calculations revealed that these data [11, 12] are consistent with the cytoplasm continuing to expand at the slower ‘meristem’ rate throughout the EZ, with a vacuolar expansion rate approximately five times higher (SI Appendix, section 7); such expansion results in a vacuolar volume fraction of 90 % once cells enter the mature zone.

We modelled GA_4_ metabolism within every cell. GA_4_ biosynthesis involves a series of oxidation steps, converting the precursor, geranylgeranyldiphosphate, to bioactive GA_4_ [43, 17]. GA_4_ biosynthesis has been shown to be predominantly regulated at the later steps of this pathway, whereby GA_12_ is converted to bioactive GA_4_ [13]. Thus, in every cell, we modelled the network (Fig 1B) whereby GA_12_ undergoes a series of oxidation steps mediated by GA 20-oxidase (GA20ox) to produce GA_9_, which is converted to the bioactive GA_4_ by GA 3-oxidase (GA3ox) [43]. The steps in the GA_4_ biosynthesis pathway are modelled via Michaelis-Menten kinetics for the metabolic reactions [6]; parameter values for GA20ox-mediated steps are estimated in [6] using data from [3], and parameter values for the GA3ox-mediated step are estimated in [40]. GA 2-oxidase (GA2ox) enzymes mediate catabolism, which we model via linear terms [24]. Here we consider two groupings of the GA2oxidase family: GA2oxidases which mediate catabolism of GA_9_ and GA_4_ (AtGA2ox1, AtGA2ox2, AtGA2ox3, AtGA2ox4, AtGA2ox6), and GA2oxidases which mediate the catabolism of GA_12_ (AtGA2ox7, AtGA2ox8, AtGA2ox9, AtGA2ox10) [22]. Transcriptomics data [23, 28] suggest levels of GA20ox, GA3ox and GA2ox vary across the root growth zones, and similar trends are seen in [1]. Assuming the enzyme activities correlate with the transcript levels, we prescribed enzyme rates depending the cell’s zone, further subdividing the meristem into the quiescent-centre zone (QCZ, 0 − 40 *µ*m from the QC) and division zone (DZ, between 40 − 240 *µ*m from the QC), based on the data in [23, 28] (Fig. 1C). While expression of most early GA_4_ biosynthetic enzymes extends throughout the root growth zone, the first committed step in GA_12_ synthesis, ent-copalyl diphosphate synthase (CPS), exhibits high expression close to the QC and low expression in the DZ and EZ [28], and thus we assumed that GA_12_ is locally synthesized in the QCZ. Furthermore, we assumed GA_12_ is delivered in the phloem-unloading zone (the rootward half of the EZ [29]), motivated by grafting studies that suggested that GA_12_ is the main mobile gibberellin and is transported from shoot-to-root within the phloem [26]. These studies used the *kao1kao2* line which lacks the final step in GA_12_ synthesis, and showed that grafting a wildtype shoot to a *kao1kao2* mutant root partially restored root growth compared with using a *kao1kao2* shoot (that lacks GA_12_ synthesis everywhere) [26]. These data provide evidence that root growth utilizes both locally synthesised GA_12_ and GA_12_ delivered from the shoot: our model enables us to assess the relative roles of these two GA_12_ sources.

To investigate the role of transport, for each gibberellin metabolite in the modelled pathway, we incorporated passive diffusion across the plasma membrane and tonoplast, and active transport via membrane carriers of the NPF family, considering influx carriers (NPF3.1 and NPF2.12), to be on the plasma membrane (importing from the apoplast to the cytoplasm) and an efflux carrier, NPF2.14, on the tonoplast (exporting from the cytoplasm to the vacuole) [35, 9]. We note that, in the root growth zone, these NPF carriers are localized on specific cell types (NPF3.1 in the endodermal plasma membranes [35], and NPF2.12 and NPF2.14 in the pericycle plasma membranes and tonoplasts, respectively [9]), and we therefore consider simulations with and without these transporters in our generic cell-file model. We note that SWEET transporters [18] are not modelled, as within the root, transcriptomics data suggest expression only in the stele in the late EZ and mature zone [1]. We further assumed that each gibberellin metabolite diffuses sufficiently fast that the concentrations can be considered to be spatially uniform within each cell cytoplasm and vacuole, as explained in [19].

As for other plant hormones, only protonated metabolites can passively diffuse into cells, whereas the NPF carriers transport the anionic forms [20, 9]. In the acidic apoplast, ~7% of GA_4_ is protonated and able to passively diffuse into cells, with the remaining 93% being anionic requiring influx via NPF3.1 and NPF2.12 [19]. In the cytoplasm over 99% of GA_4_ is anionic, and so efflux is predominantly via the NPF2.14 carriers for cytoplasmic export into the vacuole [19]. Transporter permeabilities are estimated from oocyte data [35, 42]. These estimates revealed that protonated GA_12_, GA_15_, and GA_9_ diffuses passively across cell membranes faster than protonated GA_24_ and GA_4_ but the corresponding anionic forms have slower NPF-mediated active transport (SI Appendix Table S2); these differences are thought to be due to the specific structure of the metabolites [42].

The model included diffusion within the apoplast [21], which provides an alternative continuous pathway for hormone diffusion [20]. The model incorporated diffusion between adjacent cells through plasmodesmata [32, 25], which our previous generic-cell-file model suggested may be a significant pathway for GA_4_ movement [19].

The above model assumptions are represented by a system of ordinary differential equations (ODEs) for the concentration of each gibberellin metabolite in every cellular compartment, as detailed in SI Appendix sections 1-5.

### Continuum approximation can reproduce the cell-based model solutions

To gain insight into the underlying mechanisms and reduce the model’s computational cost, we derived a continuum approximation of the above cell-based model. The approach enables us to upscale the discrete cell-based model to a macroscale continuum model, where, instead of tracking concentrations in individual cellular compartments, we introduce macroscale concentration variables continuously varying with distance.

We previously derived a continuum approximation of a simple GA_4_ model [19], which considered transport in a single file of growing identical cells (with no synthesis and degradation). This analysis showed that GA_4_ transport in a cell file can be represented by a continuum model, whereby GA_4_ concentration is a function of time and distance and the GA_4_ dynamics are governed by a single reaction–advection–diffusion equation.

Building on this previous analysis [19], we derived a continuum approximation of GA_4_ transport and metabolism across the three root growth zones (meristem, EZ, and MZ) (SI Appendix, sections 1-4, Fig S3). In each zone, the analysis led to a system of coupled partial differential equations, i.e. one equation for each metabolite, that contains terms representing advection, diffusion, dilution and production/degradation (Fig. 1D), providing formulae for the effective velocity, diffusivity, and dilution rate, in terms of the parameters governing the cell-scale metabolism, transport and cell-growth dynamics. These formula revealed that plasmodesmatal diffusion, apoplastic diffusion, and NPF-mediated transport contribute to the diffusion terms, whereas cell growth and spatial variance in the cell lengths and the subcellular compartmentalization lead to the advection and dilution terms. The systems of equations representing the dynamics in each zone are then coupled at the boundaries between the three zones. We note that a continuum approximation of GA_1_ dynamics in the Maize leaf growth zone was previously presented in [5], albeit with no transport between adjacent cells which resulted in a much simpler derivation and no diffusion terms.

The continuum approximation greatly simplified the computational process of simulating the model, enabling us to introduce more complex dynamics and perform more detailed parameter surveys. The continuum approximation also provided insights into the model dynamics via the analytic expressions for key effective parameters, which revealed how the cellscale processes contribute to the effective diffusivity, induced hormone velocity, and dilution rate.

### Model predictions can reproduce GA_4_ gradient observed using the nlsGPS1 sensor

The model predicts the distribution of the modelled gibberellin metabolites along the root growth zones (considering the steady-state distributions). Having predicted the GA_4_ distribution, the corresponding distribution of the nlsGPS1 emission ratio is calculated using the nonlinear relationship suggested from titration curves in [27]. Estimates for most model parameters are available in literature (SI Appendix, Tables S1-S3). However, the synthesis/delivery rates for GA_12_ and degradation rates for GA_12_, GA_9_, and GA_4_ are unknown. To identify appropriate values for these parameters, we performed a parameter survey comparing the predicted distribution of the nlsGPS1 emission ratio with experimental data (SI Appendix, section 6, Fig S4). The parameter survey suggests that the degradation rates for GA_12_, GA_9_ and GA_4_ are approximately equal, and that the synthesis/delivery rates for GA_12_ in the QCZ and the phloem unloading-zone are of a similar magnitude (SI Appendix, Table S4).

The GA_12_ locally synthesised in the QCZ is predicted to create a GA_4_ distribution with a peak within the DZ (Fig. 2A). The GA_12_ delivered in the phloem-unloading zone is predicted to generate higher GA_4_ levels, with GA_4_ levels increasing as cells reach the end of the meristem (Fig. 2B). Combining both sources of GA_12_ results in a superposition of the two GA_4_ distributions, with the phloem-delivered GA_12_ making a larger contribution (Fig. 2C). The corresponding predicted nlsGPS1 distribution agrees well with the experimental data (Fig. 2D). As expected, the model predicts that increasing either the rate GA_12_ synthesis in the QCZ or the rate of GA_12_ delivery via the phloem increases the GA_4_ levels (SI Appendix, Fig. S5); albeit, with increasing the GA_12_ delivery rate having a more substantial effect than increasing the GA_12_ synthesis rate (Fig. 2C).

**Figure 2:**
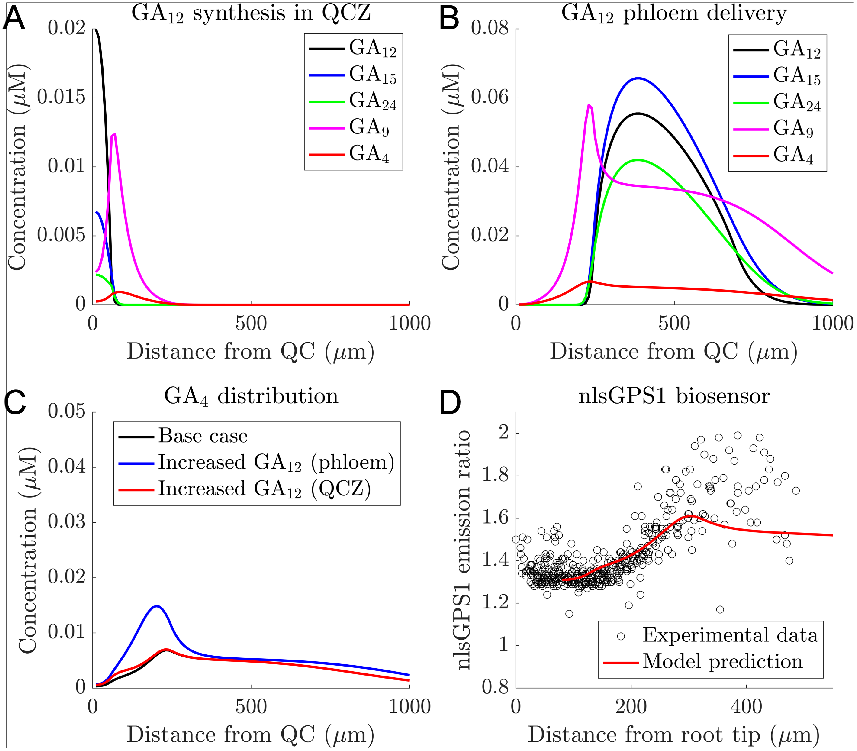
Model predictions can reproduce GA_4_ gradient observed using the nlsGPS1 sensor. (**A**,**B**) Predicted GA_12_, GA_15_, GA_24_, GA_9_, and GA_4_ distributions due to (**A**) local GA_12_ synthesis in the QCZ, and (**B**) GA_12_ delivery in the phloem-unloading zone. (**C**) Predicted GA_4_ distribution (due to both GA_12_ delivery and local synthesis), together with predictions showing the effect of doubling either the GA_12_ synthesis rate in the QCZ, or the GA_12_ delivery rate in the phloem-unloading zone. (**D**) Comparison between the predicted and observed distribution of nlsGPS1 emission ratio. Predicted sensor distribution is calculated from the base case GA_4_ distribution shown in panel C.

We used the model to evaluate the roles of the spatial distributions of both GA_12_ synthesis/delivery and enzyme levels. The model predicted that the GA_4_ distribution relies on the distribution of GA_12_ synthesis/delivery. With a uniform GA_12_ synthesis/delivery, the model does predict a GA_4_ gradient (due to the enzyme transcript distribution); however, this gradient is closer to the root tip with a wide peak within the DZ, inconsistent with the sensor data (SI Appendix, Fig. S6). In addition, the enzyme distributions are also essential to the predicted GA_4_ gradient - with uniform enzyme levels, the predicted distributions of all the gibberellin metabolites peak where GA_12_ is delivered/synthesised, with high levels of GA_4_ in the QCZ, again inconsistent with the sensor data (SI Appendix, Fig. S7).

We conclude that the model predictions capture the GA_4_ gradient observed using the nlsGPS1 biosensor, and reveal that the GA_4_ gradient is due to the combined effect of GA_12_ synthesis and delivery and spatial variations in the metabolism enzymes. The model suggests that GA_12_ delivered to the root through the phloem is an important contributor to the GA_4_ gradient.

### Enzyme inactivation in the division zone is essential to explain the GA_4_ distributions with GA20ox and GA3ox over-expression

Overexpressing GA3ox enzyme was observed to significantly increase the nlsGPS1-detected GA_4_ in the EZ and the slope of the gradient, whereas over-expressing GA20ox enzyme had little effect [28] (Fig. 3A,B). Consistent with these observations, the model predicted that overexpression of GA3ox resulted in a larger change in nlsGPS1 emission ratios compared with overexpression of GA20ox (Fig. 3E,F). However, the predicted distribution of the nlsGPS1 emission ratio disagreed with that observed, as the model predicted over-expression of GA3ox would result in a substantial increase in GA_4_ in the meristem (Fig 3B,F). This discrepancy in the meristem was also present when comparing predicted and observed distributions for lines with both GA20ox and GA3ox over-expressed (Fig. 3C,G) and lines with *ga20ox* loss-of-function (Fig. 3D,H).

**Figure 3:**
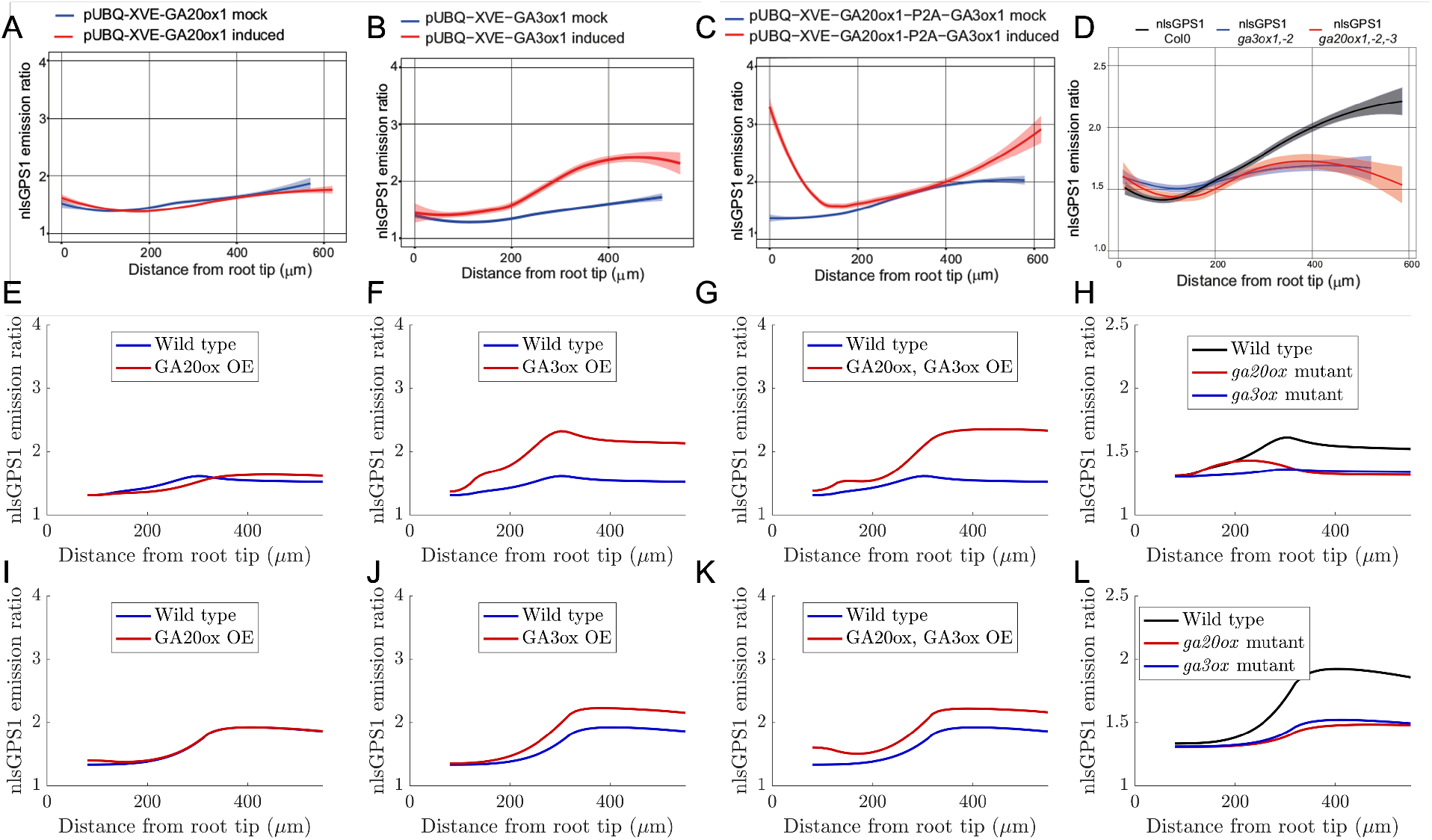
Enzyme inactivation in the division zone is essential to explain the sensor distributions in the GA20ox and GA3ox over-expression. (**A-D**) Experimental nlsGPS1 distributions reproduced from [28]. (**E-L**)Predicted nlsGPS1 emission ratio of GA_4_ for (**E**,**I**) GA20ox overexpression, (**F**,**J**) GA3ox overexpression, (**G**,**K**) both GA20ox and GA3ox overexpression, and (**H**,**L**) *ga20ox* and *ga3ox* mutants. (**E-H**) Model predictions with enzyme activities based on transcript levels (shown in Fig 1B). (**I-L**) Model predictions in which the GA20ox and GA3ox activity is set to zero in the DZ (with all other enzyme activities based on the transcript level). Model predictions with alternative assumptions are shown in Supp. Fig. S8.

Given we observed no significant increase in GA_4_ levels in the central meristematic zone during the simultaneous induction of GA20ox and GA3ox expression (Fig. 3C), we hypothesised that these biosythesis enzymes could be post-transcriptionally downregulated. Testing this hypothesis with the model, we reduced the activity of GA3ox and GA20ox either (i) throughout the meristem (SI Appendix, Fig. S8E-H, M-P) or (ii) the division zone only (Fig. 3I-L, SI Appendix Fig. S8I-L). Noting that we re-performed the parameter survey in these cases in order to identify synthesis, delivery and degradation rates for which the wild-type predictions agree with the nlsGPS1 data (SI Appendix, Fig S9, Table S4). We found that the model predictions are consistent with the data provided GA20ox and GA3ox activity are set to zero in the division zone only (Fig. 3I-L). We note that these model predictions underestimate the increase in GA_4_ levels at the QCZ in the case of double over-expression of GA20ox and GA3ox (compare Fig. 3C,K); however, they reproduce the qualitative trend. Predictions which also included zero activity in the QCZ did not agree with the data (SI Appendix, Fig. S8E-H), as they did not allow for the observed local increase of the GA_4_ levels in the QCZ in the GA20ox and GA3ox over-expression mutant (Fig. 3C). We further tested the case in which the activity of GA3ox and GA20ox are reduced to a small, but non-zero value either (i) in the division zone only (SI Appendix, Figs. S8I-L) or (ii) throughout the meristem (SI Appendix, Figs. S8M-P). We see that quantitatively the predictions scale as expected, and do not reproduce the experimental data as well as the predictions with complete enzyme inactivation.

Interestingly, the agreement between the predictions and data for wild type is also improved when we set GA20ox and GA3ox activity to be zero in the DZ (Fig. 4D). With inactive biosynthesis enzymes in the DZ, GA_12_ is able to diffuse further. As a result, the GA_12_ synthesised in the QCZ leads to GA_4_ synthesis in the EZ, and hence, both local GA_12_ synthesis in the QCZ and GA_12_ delivery via the phloem contribute to the GA_4_ gradient (Fig. 4A-C). However, the model predicts that GA_12_ delivery still makes the larger contribution to the GA_4_ levels, and increasing the rate of GA_12_ phloem delivery is predicted to have a more substantial effect than increasing the rate of local GA_12_ synthesis in the QCZ (Fig. 4C). With inactive biosynthesis enzymes in the DZ, the model predicts that the spatial distribution of GA_12_ synthesis/delivery has a lesser effect on the GA_4_ gradient: with uniform GA_12_ synthesis/delivery, the model predicts a GA_4_ gradient which is close to that observed via the nlsGPS1 sensor (SI Appendix, Fig. S10).

**Figure 4:**
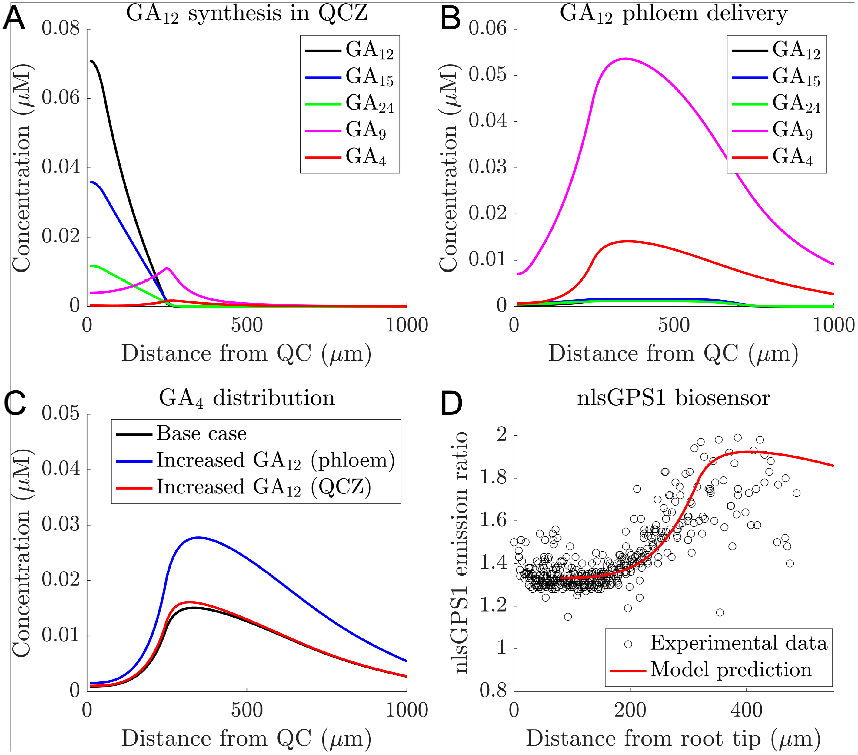
Model predictions with inactive GA20ox and GA3ox in the DZ reproduce GA gradient observed using the nlsGPS1 sensor. (**A**,**B**) Predicted GA_12_, GA_15_, GA_24_, GA_9_, and GA_4_ concentration distributions due to (**A**) GA_12_ synthesis in the QCZ and (**B**) GA_12_ delivery the phloem-unloading zone. (**C**) Predicted GA_4_ distribution (due to both GA_12_ delivery and local synthesis) showing the effect of increasing either the GA_12_ synthesis rate in the QCZ, or the GA_12_ delivery rate in the phloem-unloading zone. (**D**) Comparison between predicted and observed nlsGPS1 GA sensor distribution. Predicted sensor distribution is calculated from the base case GA_4_ distribution shown in panel C.

We conclude that there is post-transcriptional regulation reducing the activity of GA20ox and GA3ox in the dividing cells. Such enzyme inactivation is essential for reproducing the observed GA_4_ distribution in the GA20ox and GA3ox OE lines, and improves agreement between predictions and data in the wildtype, *ga3ox* and *ga20ox* lines. Furthermore, the model reinforces previous suggestions that GA3ox mediate the limiting step in GA_4_ biosynthesis in the root growth zones [28].

### Plasmodesmatal diffusion refines the GA_4_ gradient

Plasmodesmatal diffusion has been shown to have a significant effect on the distributions of other hormones [25], although its role in GA_4_ distribution has not been explored. The model predicts that plasmodesmatal diffusion has a significant effect on the GA_4_ distribution: reducing/increasing the plasmodesmatal permeability increases/decreases the slope of the GA_4_ gradient (Fig. 5A, SI Appendix, Fig. S11). With lower plasmodesmatal diffusion, GA_12_ is unable to diffuse away from its synthesis/delivery regions (SI Appendix, Fig. S11A), and the downstream metabolites are also unable to diffuse leading to a steep GA_4_ gradient as cells enter the EZ (Fig. 5A). In contrast, with higher plasmodesmatal diffusion, the model predicts that the metabolites diffuse easily into the DZ, and the slope of the GA_4_ gradient is substantially reduced (Fig. 5A).

**Figure 5:**
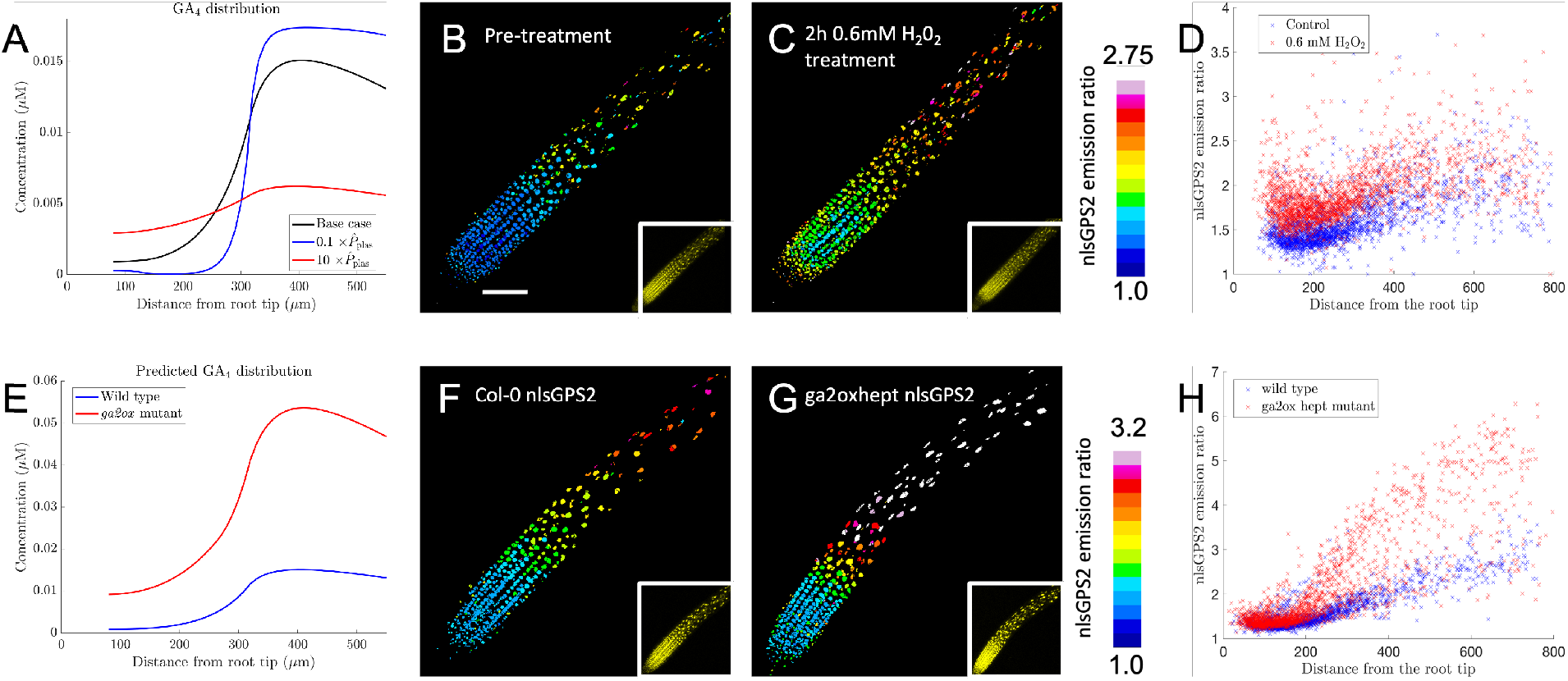
The GA_4_ gradient depends on plasmodesmatal diffusion and catabolism. (**A**) Predicted effect of the plasmodesmatal permeability, 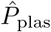, on the GA_4_ distribution. (**B**,**C**) Representative images of nlsGPS2 emission ratios and YFP fluorescence (inset) for pre-treated (**B**) and 2h 0.6mM H_2_O_2_ treated(C)roots. Treatment used to show the effect of increasing plasmodesmata permeability. Scale bar = 100 *µ*m applies to all images. (**D**) Quantification of nlsGPS2 emission ratio data for pre-treated and 2h 0.6mM H_2_O_2_ treated roots. Data quantified from images shown in panels B and C, and SI Appendix Fig S12. (**E**) Predicted GA_4_ distribution for the *ga2ox* heptuple mutant (with reduced degradation of GA_12_, GA_9_ and GA_4_), (**F**,**G**) Representative images of nlsGPS2 emission ratios and YFP fluorescence (inset) for Col0 (**F**) and *ga2oxhept* mutant (**G**) roots. (**H**) Quantification of nlsGPS2 emission ratio data for the *ga2ox* hept mutant. Data quantified from images shown in panels F and G, and SI Appendix Fig S14.

To test the model predictions that plasmodesmatal diffusion affects GA_4_ distribution, we experimentally perturbed plasmodesmatal permeability using a H_2_O_2_ treatment, choosing a treatment time of 2 h and concentration of 0.6 mM which has previously been shown to increase plasmodesmatal permeabilities within the root [32]. Experiments were performed using the GPS2 FRET biosensor, which exhibits improved orthogonality and reversibiliy that is ideal for examining GA_4_ levels and in conjunction with growth phenotypes [16]. Applying a 0.6 mM H_2_O_2_ treatment led to a shallow GA_4_ gradient (Fig. 5B-D, SI Appendix Fig. S12) in agreement with the model predictions (Fig. 5A). Interestingly, increasing plasmodesmatal permeability was also observed to lead to a small increase in GA_4_ levels within the EZ (Fig. 5D). Given phloem unloading also occurs through plasmodesmata [29], we suggest the observed increase in GA_4_ is likely due to increased phloem delivery.

We conclude that appropriate plasmodesmatal diffusion is essential for shaping the GA_4_ gradient if the plasmodesmatal diffusion is too high, the gradient will be eliminated, whereas if the diffusion is too low, the gradient would be much steeper than that observed.

### Dilution and NPF-mediated transport have limited effect on the predicted GA_4_ distribution

Since the cells grow, this naturally introduces dilution along the root, which would reduce concentrations. The model predicts that reducing dilution (by reducing the value of the cells’ RER) has a little effect on GA_4_ levels in the meristem and a small increase in GA_4_ levels in the EZ where the cell elongation is most rapid (SI Appendix, Fig. S13A).

Considering the NPF-mediated membrane transport, the model predicts a small increase in GA_4_ in the EZ with vacuolar import removed (as in a *npf2*.*14* loss-of-function mutant), and little change in the meristem (SI Appendix, Fig. S13B). Since cell elongation in the EZ is predominantly due to vacuolar expansion, removing vacuolar import reduces dilution, explaining why these predictions are similar to those with reduced RER. The model predicts that removing the NPF cytoplasmic importers (such as in a *npf3*.*1* mutant) has no noticeable effect on the GA_4_ gradient (SI Appendix, Fig. S13B).

We conclude that, while dilution and NPF2.14mediated vacuolar import are predicted to have a small effect on the GA_4_ level in the EZ, plasmodesmatal diffusion is the dominant transport mechanism that controls the GA_4_ gradient.

### Catabolism has significant effect on GA_4_ levels

We explored the effect of catabolism on GA_4_ distribution. Considering the heptuple *ga2ox* loss-of-function mutant [16], which has reduced catabolism of GA_12_, GA_9_ and GA_4_, the model predicts GA_4_ levels are higher than the wild type throughout the growth zones (Fig. 5E). To test this prediction, we examined nlsGPS2 biosensor emission ratios in the *ga2ox* heptuple mutant [16]. The nlsGPS2 data revealed higher GA_4_ levels in the *ga2ox* heptuple mutant providing support for the model prediction (Fig. 5F-H, Supp. Fig. S14).

Interestingly, the nlsGPS2 data showed that the loss of degradation in the heptuple *ga2ox* mutant had a greater effect on the GA_4_ level in the EZ than in the meristem, suggesting that the GA2ox are particularly functional in the EZ. We note that the model predicts that reduced degradation also leads to higher GA_4_ in the meristem, in contrast to the *ga2oxhept* data (compare Fig. 5E,H); this discrepancy could suggest there are other catabolic enzymes, for example, the ELAs, which are mediating GA_4_ degradation in the meristem.

Increasing the individual degradation rates (associated with GA_12_, GA_9_ or GA_4_) is predicted to lower the GA_4_ level (SI Appendix, Fig. S15). Consistent with this prediction, Kubalova et al. showed overexpression of GA2ox8 in the nlsGPS2 background resulted in lower GA_4_ levels (*Kubalova et al 2024 companion manuscript*). Increasing either the GA_4_ or GA_9_ degradation rates is predicted to result in a more substantial reduction of GA_4_ levels than increasing the GA_12_ degradation rate (SI Appendix, Fig. S15), although we note that increasing GA_12_ degradation may also reduce GA_12_ delivery from the shoot, which would further reduce the predicted GA_4_ levels.

We conclude that GA_4_ catabolism plays a key role in controlling GA_4_ levels and is essential to the GA_4_ gradient.

## Discussion

GA_4_’s role in plant growth regulation is well-established [36]. Our previous studies discovered a GA_4_ gradient in the Arabidopsis root growth zones that appears to underlie root cell elongation [27, 28]. By developing a detailed multiscale mathematical model, we now reveal how cell-scale processes combine to create this GA_4_ gradient, via the combination of GA_12_ synthesis and delivery, spatial variations in biosynthesis enzyme activity, plasmodesmatal diffusion and catabolism.

The model revealed that inactivation of the GA20ox and GA3ox biosynthesis enzymes in the DZ is essential to the GA_4_ gradient. We had previously observed that over-expressing the GA20ox and GA3ox enzymes did not increase GA_4_ levels in the DZ [28]; comparing model predictions with these previous data revealed that these enzymes are inactive in the DZ. Furthermore, we found that enzyme inactivation refined the predicted wild-type gradient, leading to better agreement between predictions and data. This finding is aligned with that in Maize leaves that suggested post-transcriptional regulation to be important in regulating GA_1_ levels in response to drought and cold [5], where GA_1_ plays a similar role in regulating meristem size.

Our study revealed that plasmodesmatal diffusion antagonizes the GA_4_ gradient. Plasmodesmatal diffusion has been shown to affect distributions of auxin [25], but its influence on other hormone distributions was yet to be evaluated. Given plasmodesmatal permeabilities are rapidly regulated by environmental conditions [34, 8], a resulting modification to the GA_4_ gradient may provide an additional mechanism for environmental regulation of root growth.

In contrast to the role of plasmodesmatal diffusion, the model predicted that the NPF-mediated active influx across the cell membranes had limited effect on the GA_4_ distribution. Given cytoplasmic GA_4_ is predominantly anionic and unable to cross membranes via passive diffusion [20], we predicted that the apoplastic GA_4_ concentrations are low and there is little GA_4_ available for influx via the plasma membrane carriers (NPF3.1, NPF2.12). Should GA_4_ efflux carriers be identified in the future, we anticipate that, where efflux occurs, apoplastic GA_4_ levels would be higher and influx carriers would then be predicted to contribute to the GA_4_ distribution. Plasma membrane import may also have an important role with exogenous gibberellin, for example NPF transporters were shown to affect the distribution of fluorescently-tagged gibberellin between different tissues in the root cross-section [35, 9].

The model predictions suggest that phloem delivery of GA_12_ provides an important source of precursor to create high GA_4_ levels as cell’s enter the elongation zone. However, this does not rule out the possibility that local GA_12_ synthesis also plays an important role: sensor emission ratios suggest a small peak in GA_4_ levels close to the QC, which the model simulations suggest is derived from locally synthesised GA_12_ in this region. The strong GA_4_ in the QCZ with over-expression of both GA3ox and GA20ox also indicates a local pool of GA_12_ [28]. A recent study that found inducing GA_4_ degradation specifically in the QCZ reduced meristem length, whereas inducing GA_4_ degradation in the shootward-meristem and EZ affected elongated cell length but not meristem length [7]. If the QCZ and EZ GA_4_ pools do indeed have distinct functions and can be regulated independently as the model suggests, roots could use GA_4_ distribution to separately influence meristem size and cell elongation.

The model explains the perturbed GA_4_ distributions observed in several characterized mutants. We predicted GA_4_ distributions for mutation and over-expression of GA2ox enzymes that have been corroborated experimentally in nlsGPS2 lines in *ga2ox* heptuple and GA2ox8 overexpression backgrounds (Fig 5, and Kubalova et al 2024, companion manuscript). The increased and reduced growth phenotypes of *ga2ox* heptuple and GA2ox8 overexpression lines evidence the quantitative relationship between cellular GA distribution and localized root growth (Kubalova 2024 companion manuscript).

There have been numerous studies of plant hormone distributions focused on the hormone auxin, investigating how the auxin distributions are created, and how they mediate developmental processes and environmental responses [14]. In comparison, our understanding of the distributions of other plant hormones is in relative infancy. Recently developed FRET sensors have revealed that other hormones, such as GA and ABA, also have dynamic spatial distributions, that are affected by genetic mutation and environmental conditions [27, 28, 30, 16, 39, 10]. As in the auxin field [33], computational models, like that developed here, will be vital to explain and understand these hormone distributions and how they arise through processes at the cellular scale. We anticipate the model developed here will provide a basis for future modelling studies, incorporating additional processes such as hormone cross-talk or multiple tissue layers.

We conclude that GA_4_ distributes like a classical morphogen, as proposed by Wolpert [41], with localized synthesis combining with diffusion and degradation to create a spatial gradient. At the same time, GA_4_ does not alone control the strict longitudinal patterning of QCZ, meristem and EZ of the Arabidopsis root. Interestingly, the results presented in the companion manuscript show that auxin regulation of GA_4_ catabolism mediates growth responses (Kubalova et al, 2024). These findings could point to the GA_4_ gradient being a hub for other hormone signals, with regulation of our model inputs enabling other signals to regulate growth. The clear roles of GA_4_ in contributing to the root growth parameters suggest that other developmental pattern generators likely direct the GA_4_ enzymes and transport regimes uncovered here.

## Materials and Methods

### Mathematical modelling

Full details of the derivation of the model equations are provided in the SI Appendix sections 1-4. Where available, model parameter estimates are obtained from the literature (see Tables S1-3), and the remaining five parameter values are estimated by comparing predictions with nlsGPS1 data (see SI Appendix section 5, Table S4). Simulations were performed using matlab.

### Plant material and growth conditions

WT and mutant lines used in this study were *A. thaliana* ecotype Columbia 0 (Col-0). Seeds were chlorine-gas-sterilized and plated on 1/2 Murashige and Skoog (MS) basal medium (Duchefa, Cat No. M0221) with 0.025% 2-morpholinoethanesulfonic acid monohydrate (MES), pH 5.7, and 1.2% agar (1/2 MS solid medium, Sigma). After stratification in the dark at 4 °C for 3 d, plates were placed in a growth chamber with long-day growth conditions (120 *µ*mol m^*−*2^ s^*−*1^ white light, 22 °C for 16 h; 0 *µ*mol m^*−*2^ s^*−*1^, 18 °C for 8 h). Lines used in this study nlsGPS2 for both Col-0 and *ga2oxhept* [16].

### Confocal imaging and treatments

For steady state experiments, samples were mounted in liquid 1/4 × MS medium (1/4 × MS salts, 0.025% MES, and pH 5.7) with coverslips and imaged. For the 0.6mM H_2_0_2_ treatment, the standard medium beneath the coverslip was exchanged with the medium containing H_2_0_2_ solution. The H_2_0_2_ medium was prepared immediately before adding to the sample. A control buffer exchange was also carried out. The samples were then re-imaged after 2 hours. Confocal images were acquired with a format of 1024 × 512 pixels and resolution of 12 bit on an upright Leica SP8 using a 20× dry 0.70 HC PLAN APO objective. To excite Cerulean and Aphrodite, 448 nm and 514 nm lasers were used, respectively. Emission filters were 460 to 500 nm for Cerulean and 525 to 560 nm for Aphrodite. Three fluorescence channels were collected for FRET imaging: Cerulean donor excitation and emission or DxDm, Cerulean donor excitation, Aphrodite acceptor emission or DxAm, and Aphrodite acceptor excitation and emission or AxAm.

### Image processing and analysis

Imaging process and analysis were performed with FRETENATOR plugins [31]. Segmentation settings were optimized for each experiment but kept constant within each experiment. The AxAm channel was used for segmentation. For segmentation, Otsu thresholds were used, difference of Gaussian kernel size was determined empirically, and a minimum ROI size was set to 20. Distance from meristem was defined using FRETENATOR ROI labeler.

KBK and LRB are grateful to the Human Frontier Science Program (HFSP) (grant number: RGY0075/2020) for subsidising this work. LRB is grateful to the BBSRC (grant number: BB/S001190/1) for subsidising this work. JG and AMJ are grateful to the Gatsby Charitable trust (GAT3395) and European Research Council under the European Union’s Horizon 2020 research and innovation program (grant agreement 759282). We thank Eilon Shani and Hussam Nour-Eldin for helpful discussions.

## Notes

### Competing Interest Statement

The authors have declared no competing interest.

