## Supplemental text, Supplemental Figures S1-S15, Supplemental Tables S1-S4 for "A balance of metabolism and diffusion articulates a gibberellin hormone gradient in the *Arabidopsis* root"

March 24, 2025

In this Supplementary Information, we provide further details on the mathematical modelling.

We begin by describing the cell-based model for transport of a single hormone in a generic file of growing and dividing cells with constant hormone synthesis and degradation rates. We then use the results in [14] to derive an upscaled continuum model for the hormone concentration. In Section 1, we present the cell-based and the upscaled continuum models incorporating the hormone transport dynamics. In Section 2, we discuss the root growth dynamics and the corresponding assumptions used in the model. In Section 3, we verify that the continuum approximation recapitulates the output of the cell-based model using the growth dynamics described in Section 2. In Section 4, we describe model of the gibberellin metabolic pathway (as shown in Fig. 1B in the main text). In Section 5, we describe the full model that incorporates transport dynamics, growth, and the metabolic pathway, detailed in Sections 1–4. In Section 6, we describe how parameter estimates were obtained: the majority of the model parameter values are obtained from the literature (as listed in Tables 1-3); however, five parameter values are unavailable and were estimated by comparing model predictions with the wildtype nlsGPS1 data, as detailed in section 6 and given in Table 4.

#### 1 Modelling hormone transport via a continuum approximation of a cell-based model

Before presenting the full model, we first describe hormone transport in a model for a single hormone to demonstrate how a cell-based transport model can be approximated by a continuum equation. We consider transport in a file of length $\hat{L}(\hat{t})$ of $N$ growing and dividing cells with apoplastic (cell wall) and subcellular (vacuolar) compartments, as shown in Supp. Fig. 2. We incorporate passive transport and facilitated transport via protein transporters located on the cytoplasmic and vacuolar membranes of the cells. We include plasmodesmatal transport between cell cytoplasms and apoplastic diffusion within the cell wall. We use the notation and model description in [14] to express the rate of change of hormone concentration in each compartment in terms of the net flux of the hormone through the relevant boundaries. Further, we assume that hormone is produced at a rate $\hat{\sigma}$ and degraded at a rate $\hat{\beta}$ within each cytoplasm (noting that these terms will be replaced in the full model described in Section 5). Initially, there is no hormone in the cells. The hormone dynamics can be modelled via a system of coupled ordinary differential equations for the hormone concentrations in the cytoplasm ($\hat{c}_i$), vacuole ($\hat{v}_i$), and adjacent apoplast compartments ($\hat{f}_i, \hat{g}_i$, and $\hat{h}_i$), where, as in [14], we assume that the hormone concentrations are constant within each compartment:

$$\hat{w}\frac{\mathrm{d}}{\mathrm{d}\hat{t}}\left((1-\phi_i)\hat{l}_i\hat{c}_i\right) = \hat{w}(\hat{J}_{fci} - \hat{J}_{cfi} + \hat{J}_{cci} - \hat{J}_{cc(i-1)}) - \hat{l}_i\hat{J}_{chi} - 2\sqrt{\phi_i}(\hat{l}_i + \hat{w})\hat{J}_{cvi} + \hat{w}(1-\phi_i)\hat{l}_i(\hat{\sigma} - \hat{\beta}\hat{c}_i), \tag{1a}$$

$$\hat{w}\frac{\mathrm{d}}{\mathrm{d}\hat{t}}\left(\phi_i\hat{l}_i\hat{v}_i\right) = 2\sqrt{\phi_i}(\hat{l}_i + \hat{w})\hat{J}_{cvi}, \tag{1b}$$

$$\hat{a}\hat{w}\frac{\mathrm{d}\hat{f}_i}{\mathrm{d}\hat{t}} = \hat{w}(\hat{J}_{cfi} - \hat{J}_{fc(i+1)}) - \hat{a}\hat{J}_{fgi}, \tag{1c}$$

$$\hat{a}^2\frac{\mathrm{d}\hat{g}_i}{\mathrm{d}\hat{t}} = \hat{a}(\hat{J}_{hgi} - \hat{J}_{gh(i+1)} + \hat{J}_{fgi}), \tag{1d}$$

$$\hat{a}\frac{\mathrm{d}}{\mathrm{d}\hat{t}}\left(\hat{l}_i\hat{h}_i\right) = \hat{l}_i\hat{J}_{chi} + \hat{a}(\hat{J}_{ghi} - \hat{J}_{hgi}), \tag{1e}$$

which hold for $2 \leq i \leq N$, where $\hat{w}$ is the width of the cells, $\hat{a}$ is the thickness of the apoplastic compartments, $\hat{l}_i$ are the cell lengths, $\phi_i$ are the vacuolar area fractions in each cell, and $\hat{J}_{xyi}$ is the flux from compartment $x$ to compartment $y$ in cell $i$, i.e.,

$$\hat{J}_{cfi} = \hat{P}_{\mathrm{ca}}\hat{c}_i - \hat{P}_{\mathrm{ac}}\hat{f}_i, \tag{2a}$$

$$\hat{J}_{cvi} = \hat{P}_{\mathrm{cv}}\hat{c}_i - \hat{P}_{\mathrm{vc}}\hat{v}_i, \tag{2b}$$

$$\hat{J}_{fci} = \hat{P}_{\mathrm{ac}}\hat{f}_{i-1} - \hat{P}_{\mathrm{ca}}\hat{c}_i, \tag{2c}$$

$$\hat{J}_{chi} = \hat{P}_{\mathrm{ca}}\hat{c}_i - \hat{P}_{\mathrm{ac}}\hat{h}_i, \tag{2d}$$

$$\hat{J}_{fgi} = \frac{2\hat{D}_{\mathrm{apo}}}{\hat{w} + \hat{a}}(\hat{f}_i - \hat{g}_i), \tag{2e}$$

$$\hat{J}_{ghi} = \frac{2\hat{D}_{\mathrm{apo}}}{\hat{l}_i + \hat{a}}(\hat{g}_{i-1} - \hat{h}_i), \tag{2f}$$

$$\hat{J}_{hgi} = \frac{2\hat{D}_{\mathrm{apo}}}{\hat{l}_i + \hat{a}}(\hat{h}_i - \hat{g}_i), \tag{2g}$$

$$\hat{J}_{cci} = \hat{P}_{\mathrm{plas}}(\hat{c}_{i+1} - \hat{c}_i). \tag{2h}$$

Descriptions of the remaining parameters can be found in Supplementary Tables 1 and 2, and further explanation is provided in [14].

We assume the hormone cannot leave the file at either end, and thus, the governing equations (1) are subject to the following boundary conditions

$$\hat{c}_1 = \hat{c}_2, \qquad \hat{h}_1 = \hat{h}_2, \tag{3a}$$

$$\hat{c}_{N+1} = \hat{c}_{N-1}, \qquad \hat{h}_{N+1} = \hat{h}_{N-1}, \tag{3b}$$

and initial conditions

$$\hat{c}_i = \hat{v}_i = \hat{f}_i = \hat{g}_i = \hat{h}_i = 0 \qquad \text{at} \qquad \hat{t} = 0 \qquad \text{for } 2 \leq i \leq N. \tag{4}$$

We note that, in contrast to [14], here, we have additional production and degradation terms in equation (1a), whereas (3) are no-flux (Neumann), rather than a Dirichlet condition.

Following the analysis in [14], we upscale (1)–(4) to derive a continuum description of the system, where all variables (hormone concentrations, cell lengths, and vacuolar fractions) become functions of space, $\hat{x}$, along the file and time, $\hat{t}$, e.g., $\hat{c}_i \approx \hat{C}(\hat{x}, \hat{t})$, $\hat{l}_i(\hat{t}) \approx \hat{l}(\hat{x}, \hat{t})$, etc. This approach greatly simplifies the computational challenges associated with solving for the discrete model in every cell accounting for cell growth and division. Additionally, through the derivation of effective upscaled parameters, such as effective diffusivity and advective velocity, it provides insight into what the effect of each parameter on the system behaviour is.

Exploiting the fact that the cell lengths are much less than the tissue length, it can be shown that the hormone concentrations in the apoplastic and vacuolar compartments are approximately proportional to the cytoplasmic concentration (see [14]), i.e.,

$$\hat{f} = \hat{g} = \hat{h} = \mathcal{P}_{\mathrm{a}} \hat{C}, \qquad \hat{v} = \mathcal{P}_{\mathrm{v}} \hat{C}, \tag{5}$$

where

$$\mathcal{P}_{\mathrm{a}} = \hat{P}_{\mathrm{ca}}/\hat{P}_{\mathrm{ac}}, \qquad \mathcal{P}_{\mathrm{v}} = \hat{P}_{\mathrm{cv}}/\hat{P}_{\mathrm{vc}}. \tag{6}$$

With this, following the derivation in [14], we can show that the discrete model (1) can be approximated by a continuum reaction–advection–diffusion equation for the upscaled cytoplasmic concentration $\hat{C}(\hat{x}, \hat{t})$. This takes the following form

$$\frac{\partial \hat{C}}{\partial \hat{t}} + \frac{\partial}{\partial \hat{x}} \left( \hat{U}_{\mathrm{eff}}(\hat{x}, \hat{t}) \hat{C} \right) = \frac{\partial}{\partial \hat{x}} \left( \hat{D}_{\mathrm{eff}}(\hat{x}, \hat{t}) \frac{\partial \hat{C}}{\partial \hat{x}} \right) - \hat{Q}_{\mathrm{eff}}(\hat{x}, \hat{t}) \hat{C} + \frac{(1-\phi)\hat{l}(\hat{\sigma} - \hat{\beta}\hat{C})}{\hat{V}}, \tag{7}$$

where

$$\hat{U}_{\mathrm{eff}}(\hat{x}, \hat{t}) = \hat{u} + \frac{\hat{K}(\hat{l} + \hat{a}) + \hat{M}}{\hat{V}} \frac{\partial \hat{l}}{\partial \hat{x}} - \frac{(\hat{K}(\hat{l} + \hat{a}) + \hat{M})(\hat{l} + \hat{a})}{\hat{V}^2} \frac{\partial \hat{V}}{\partial \hat{x}}, \tag{8a}$$

$$\hat{D}_{\mathrm{eff}}(\hat{x}, \hat{t}) = \frac{(\hat{K}(\hat{l} + \hat{a}) + \hat{M})(\hat{l} + \hat{a})}{\hat{V}}, \tag{8b}$$

$$\hat{Q}_{\mathrm{eff}}(\hat{x}, \hat{t}) = \frac{\hat{u}}{\hat{V}} \frac{\partial \hat{V}}{\partial \hat{x}} - \frac{\partial \hat{U}_{\mathrm{eff}}}{\partial \hat{x}} + \frac{1}{\hat{V}} \frac{\partial \hat{V}}{\partial \hat{t}} = \frac{1}{\hat{V}} \frac{\mathrm{d}\hat{V}}{\mathrm{d}\hat{t}} - \frac{\partial \hat{U}_{\mathrm{eff}}}{\partial \hat{x}}, \tag{8c}$$

and, for convenience, we have defined

$$\begin{gathered} \hat{K} = \hat{w}\hat{P}_{\mathrm{ca}}/2 + \hat{w}\hat{P}_{\mathrm{plas}}, \quad \hat{M} = \hat{D}_{\mathrm{apo}}\hat{a}\mathcal{P}_{\mathrm{a}}, \\ \hat{V} = (1-\phi)\hat{w}\hat{l} + \phi\hat{w}\hat{l}\mathcal{P}_{\mathrm{v}} + \hat{a}(\hat{w} + \hat{a} + \hat{l})\mathcal{P}_{\mathrm{a}}, \quad \mathcal{P}_{\mathrm{v}} = \hat{P}_{\mathrm{cv}}/\hat{P}_{\mathrm{vc}}, \quad \mathcal{P}_{\mathrm{a}} = \hat{P}_{\mathrm{ca}}/\hat{P}_{\mathrm{ac}}, \end{gathered} \tag{9}$$

where $\hat{u}(\hat{x}, \hat{t})$ is the advective velocity due to growth calculated from the growth rate, as in [14]. From (3) and (4), this is subject to

$$\frac{\partial \hat{C}}{\partial \hat{x}} = 0 \quad \text{at} \quad \hat{x} = 0, \tag{10a}$$

$$\frac{\partial \hat{C}}{\partial \hat{x}} = 0 \quad \text{at} \quad \hat{x}(\hat{t}) = \hat{L}(\hat{t}), \tag{10b}$$

$$\hat{C} = 0 \quad \text{at} \quad \hat{t} = 0. \tag{10c}$$

(7) subject to (10) is equivalent to the corresponding equation derived in [14], albeit with the addition of the final term in (7) that represents synthesis and degradation.

#### 2 Modelling cell growth and division within the *Arabidopsis* root

A cell file in the *Arabidopsis* root comprises three different zones with their own specific cell dynamics. The first zone, adjacent to the root tip, is the meristem, where cells rapidly divide, and the cell elongation rate is slow. We assume every cell in the meristem divides once it approximately doubles in length. The next zone, adjacent to the meristem, is the elongation zone, where cells do not divide but grow rapidly. Here, cells increase their length almost 10-fold [9]. The third zone, adjacent to the elongation zone, is the maturation zone, where cells have attained their final length and do not grow, nor divide. In the model, we take the maturation zone to extend until $\hat{x} = \hat{L}_{\text{root}}$. We assume that the metabolite concentrations and their fluxes are continuous across the boundaries between the zones.

We focus on mature seedlings, in which the sizes of the meristem and the elongation zone are stable, and denoted by $\hat{L}_{\text{meri}}$ and $\hat{L}_{\text{ez}}$, respectively. Thus, we assume the cell lengths evolve at a quasi-steady state, i.e., there is approximately the same number of cells in each zone, and the cell lengths and vacuolar fractions are only functions of distance away from the root tip. Using data in [25], we assume that the cell elongation rates in the meristem and the elongation zone are constant and equal to $\hat{\text{RER}}_{\text{meri}}$ and $\hat{\text{RER}}_{\text{ez}}$, respectively. Thus, for every cell $i$, its length evolves according to

$$\frac{\mathrm{d}\hat{l}_i}{\mathrm{d}\hat{t}} = \hat{\text{RER}}_{\text{zone}} \hat{l}_i, \tag{11}$$

where "zone" refers to either the meristem (meri) or the elongation zone (ez). Within the meristem, cells undergo division at discrete times, with the time between two successive cell divisions given by

$$\hat{T}_{\text{d}} = \frac{\ln(2)}{\hat{\text{RER}}_{\text{meri}}}, \tag{12}$$

see [10].

In the meristem, cell growth is due to both cytoplasmic and vacuolar expansion [7]. To maintain a constant vacuole volume fraction throughout the meristem, $\phi_{\text{meri}}$ (as observed in [7]), we must assume that the cytoplasm and the vacuole expand at the same rate, $\hat{\text{RER}}_{\text{meri}}$. In the elongation zone, previous models [2, 20] had assumed that cell elongation is entirely due to vacuolar expansion; however, new data [7, 8] contradicts this previous assumption and reveals that the cytoplasm continues to expand slowly in the elongation zone. Calculations detailed in section 7 below show that these new data [7, 8] are consistent with the model assumptions that the cytoplasm expands slowly at the cell elongation rate in the meristem, while the vacuole expands more rapidly to enable the cells to elongate at a rate of $\hat{\mathrm{RER}}_{\mathrm{ez}}$ (larger than $\hat{\mathrm{RER}}_{\mathrm{meri}}$). Using (36) and (37), we obtain the following governing equation for $\phi_i$ in the EZ

$$\frac{1}{1-\phi_i}\frac{\mathrm{d}\phi_i}{\mathrm{d}\hat{t}} = \hat{\mathrm{RER}}_{\mathrm{ez}} - \hat{\mathrm{RER}}_{\mathrm{meri}}. \tag{13}$$

In the mature zone, there is no cell elongation and therefore we assume the vacuolar volume fraction is constant, denoted $\phi_{\mathrm{mat}}$.

We now have enough information to solve the discrete model (1)–(4) with (11) and (13) to predict the distribution of a single gibberellin metabolite.

Having described the growth assumptions in terms of the cell-based model, we now need to determine the corresponding continuum growth model by deriving corresponding formulae for the average continuum cell lengths, $\hat{l}(\hat{x},\hat{t})$, vacuolar fractions, $\phi(\hat{x},\hat{t})$, and cell velocity, $\hat{u}(\hat{x},\hat{t})$.

In the meristem, data [7, 8, 20, 25] suggest that the average cell lengths and the vacuolar fractions are approximately spatially constant, denoted by $\hat{l}_{\mathrm{meri}}$ and $\phi_{\mathrm{meri}}$, respectively. Due to the continual cell division, we take $\hat{l}_{\mathrm{meri}}$ to be the average cell length during one cell cycle, i.e.,

$$\hat{l}_{\mathrm{meri}} = \frac{1}{\hat{T}_{\mathrm{d}}}\int_0^{\hat{T}_{\mathrm{d}}} \hat{l}_i \,\mathrm{d}\hat{t}, \tag{14}$$

where $\hat{T}_d$ is the time between two successive divisions, ((12)). We can find the relationship between the initial cell length, $l_0$, (immediately after the division event) and the average cell length, $\hat{l}_{\mathrm{meri}}$, by substituting (11) and (12) into (14), noting that the quasi-steady state assumption requires the cell length to double between two successive divisions; this gives us

$$\hat{l}_{\mathrm{meri}} = \frac{\hat{l}_0}{\ln(2)}. \tag{15}$$

With a value of $\hat{l}_0 = 6\,\mu\mathrm{m}$ from [9], we calculate the average cell length to be $\hat{l}_{\mathrm{meri}} = 8.7\mu\mathrm{m}$, which is consistent with the distributions of cell lengths shown in [9]. Given that the relative elongation rates are taken to be constant within each zone, in the quasi-steady frame of reference, the cell lengths follow a linear profile in the elongation zone with respect to distance along the root, using (11) as explained in [20]. In particular, the relation

$$\frac{\mathrm{d}\hat{l}}{\mathrm{d}\hat{x}} = \hat{c}\hat{\mathrm{RER}}_{\mathrm{ez}} \tag{16}$$

holds, where the constant

$$\hat{c} = \frac{\hat{l}_{\mathrm{meri}}}{\hat{\mathrm{RER}}_{\mathrm{meri}}\hat{L}_{\mathrm{meri}}} \tag{17}$$

is the average time between successive cells leaving the meristem, and hence entering/leaving the EZ (see [10, 26, 13, 5, 20]).

In the maturation zone, the average cell lengths and the vacuolar fractions are constant, denoted by $\hat{l}_{\mathrm{mat}}$ and $\phi_{\mathrm{mat}}$ respectively, which is consistent with data [4, 25]. By integrating (16), we see that the average cell length in the mature zone can be calculated via

$$\hat{l}_{\mathrm{mat}} = \hat{l}_{\mathrm{meri}} + \hat{c}\hat{\mathrm{RER}}_{\mathrm{ez}}\hat{L}_{\mathrm{ez}}. \tag{18}$$

The velocity is calculated using the elongation rates, as explained in [14], namely, using a mass conservation law for the cell density and expressing the gradient of the velocity as the RER in the corresponding zone. Therefore, we let

$$\hat{l}(\hat{x}) = \begin{cases} \hat{l}_{\text{meri}} & \text{if } \hat{x} \leq \hat{L}_{\text{meri}} \\ \hat{l}_{\text{meri}} + (\hat{x} - \hat{L}_{\text{meri}})(\hat{l}_{\text{mat}} - \hat{l}_{\text{meri}})/\hat{L}_{\text{ez}} & \text{if } \hat{L}_{\text{meri}} < \hat{x} \leq \hat{L}_{\text{meri}} + \hat{L}_{\text{ez}} \\ \hat{l}_{\text{mat}} & \text{if } \hat{x} > \hat{L}_{\text{meri}} + \hat{L}_{\text{ez}} \end{cases},$$

$$\phi(\hat{x}) = \begin{cases} \phi_{\text{meri}} & \text{if } \hat{x} \leq \hat{L}_{\text{meri}} \\ \phi_{\text{meri}} + (\hat{x} - \hat{L}_{\text{meri}})(\phi_{\text{mat}} - \phi_{\text{meri}})/\hat{L}_{\text{ez}} & \text{if } \hat{L}_{\text{meri}} < \hat{x} \leq \hat{L}_{\text{meri}} + \hat{L}_{\text{ez}} \\ \phi_{\text{mat}} & \text{if } \hat{x} > \hat{L}_{\text{meri}} + \hat{L}_{\text{ez}} \end{cases}, \tag{19}$$

$$\hat{u}(\hat{x}) = \begin{cases} \hat{\text{RER}}_{\text{meri}}\hat{x} & \text{if } \hat{x} \leq \hat{L}_{\text{meri}} \\ \hat{\text{RER}}_{\text{ez}}(\hat{x} - \hat{L}_{\text{meri}}) + \hat{\text{RER}}_{\text{meri}}\hat{L}_{\text{meri}} & \text{if } \hat{L}_{\text{meri}} < \hat{x} \leq \hat{L}_{\text{meri}} + \hat{L}_{\text{ez}} \\ \hat{\text{RER}}_{\text{ez}}\hat{L}_{\text{ez}} + \hat{\text{RER}}_{\text{meri}}\hat{L}_{\text{meri}} & \text{if } \hat{x} > \hat{L}_{\text{meri}} + \hat{L}_{\text{ez}} \end{cases}.$$

In Supp. Fig. 1, we show plots of the cell elongation rate, length, and velocity using equations (19) and the parameter values listed in Table 1.

### 3 Comparison between continuum and cell-based model

Before proceeding with modelling $GA_4$ metabolism, we first verified that predictions using the continuum approximation agree with those of the cell-based model using the growth dynamics detailed in Section 2. Given the continuum approximation of the transport model was already carefully verified in [14], we here focus on testing the derived growth dynamics, using a simpler transport model in which we omit the apoplast and assume $GA_4$ moves between compartments only via plasmodesmata. For concreteness, we set the spatial variations in synthesis rate using the form suggested in [20]:

$$\hat{\sigma}(x) = \sigma_{QC} + \frac{\alpha x^n}{\xi + x^n}, \tag{20}$$

with parameter values $\sigma_{QC} = 0.00005\mu$M/hr, $\alpha = 0.0006\mu$M /hr, $\xi = 125\mu$m and $n = 10$. We used a constant degradation rate, $\hat{\beta} = 0.01$/hr.

For the cell-based model, we solve the system of ODEs (1a,b) with (2h) setting $\hat{J}_{cfi} = \hat{J}_{fci} = \hat{J}_{chi} = \hat{J}_{fgi} = \hat{J}_{ghi} = \hat{J}_{hgi} = 0$. These equations are subject to boundary and initial conditions given by (3) and (4), and are coupled to the ODEs describing the growth dynamics, (11), and (13).

For the continuum model, we solve the PDE problem (7)–(10) with the growth dynamics as prescribed in (19). In solving the continuum model, we find it convenient to define the following zone-dependent formula using (9), in the meristem,

$$\frac{d\hat{V}}{d\hat{t}} = \hat{\text{RER}}_{\text{meri}}((1 - \phi_{\text{meri}})\hat{w} + \phi_{\text{meri}}\hat{w}\mathcal{P}_{\text{v}} + \hat{a}\mathcal{P}_{\text{a}})\hat{l}, \tag{21}$$

and in the elongation zone,

$$\frac{d\hat{V}}{d\hat{t}} = \hat{\text{RER}}_{\text{meri}}(1 - \phi)\hat{w}\hat{l} + (\hat{\text{RER}}_{\text{ez}} - (1 - \phi)\hat{\text{RER}}_{\text{meri}})\hat{w}\hat{l}\mathcal{P}_{\text{v}} + \hat{\text{RER}}_{\text{ez}}\hat{a}\hat{l}\mathcal{P}_{\text{a}}. \tag{22}$$

We use MATLAB *ode15s* package in both cases and run the simulations to large times to obtain the steady-state solutions. We show a comparison between the discrete and the continuum model in Supp. Fig. 3 and we see excellent agreement. Having now verified this agreement, we proceed to use the continuum formulation of the model to incorporate the metabolic pathway and transport of each gibberellin metabolite.

### 4 Modelling the $GA_4$ synthesis and degradation pathway focusing on metabolism downstream of $GA_{12}$

In order to study the biosynthesis of the bioactive form $GA_4$, we incorporate the metabolic pathway, whereby $GA_4$ is synthesised from $GA_{12}$. The $GA_{12}$ is initially converted to $GA_{15}$ under the action of the GA20ox enzymes. The GA20ox enzymes also convert $GA_{15}$ to $GA_{24}$ and convert $GA_{24}$ to $GA_9$. Under the action of the GA3ox enzymes, $GA_9$ is converted into $GA_4$. Thus, $GA_4$ synthesis is mediated via the following steps:

$$\mathrm{GA}_{12} \xrightarrow{\mathrm{GA}_{20\mathrm{ox}}} \mathrm{GA}_{15} \xrightarrow{\mathrm{GA}_{20\mathrm{ox}}} \mathrm{GA}_{24} \xrightarrow{\mathrm{GA}_{20\mathrm{ox}}} \mathrm{GA}_{9} \xrightarrow{\mathrm{GA}_{3\mathrm{ox}}} \mathrm{GA}_{4}. \tag{23}$$

Several of the GA metabolites are degraded via the GA2oxidase enzymes. Since different forms of GA2ox are involved in degradation of $GA_{12}$, $GA_9$, and $GA_4$, we denote them by GA2oxA, GA2oxB, and GA2oxC, respectively [19]. Thus, $GA_4$ degradation is mediated via the following steps:

$$\mathrm{GA}_{12} \xrightarrow{\mathrm{GA}_{2\mathrm{oxA}}} \text{degradation}, \tag{24}$$

$$\mathrm{GA}_{9} \xrightarrow{\mathrm{GA}_{2\mathrm{oxB}}} \text{degradation}, \tag{25}$$

$$\mathrm{GA}_{4} \xrightarrow{\mathrm{GA}_{2\mathrm{oxC}}} \text{degradation}. \tag{26}$$

In the model, we assume that $GA_{12}$ (i) can be produced within the quiescent centre (QCZ) region, which we assume takes up the first five cells of the meristem, and (ii) can be delivered to cells within the phloem-unloading zone, which has been shown to be approximated by the most rootward half of the elongation zone [21]. These two sources of $GA_{12}$ will be referred to from now on as the QC and phloem $GA_{12}$ pools.

### 5 Full model

In order to model the dynamics of all gibberellin metabolites, described in Section 4, we first extend the continuum model (7)–(10) from Section 1 to model transport and metabolism of each GA metabolite with the synthesis and degradation terms replaced with the corresponding terms from the metabolic pathway. Transport is accounted for through the diffusivity and the permeability values of the transporters and plasmodesmata, whereas cell growth and division is modelled as in Section 2 (see (11)). To overcome the increasing computational complexity due to the number of equations and evolving geometry involved we use the continuum approximation introduced at the end of Section 1. This leads to a system of five reaction–advection–diffusion equations for the concentration of each gibberellin metabolite. We model the metabolic reactions by assuming Michaelis-Menten kinetics for the enzyme and metabolite concentration participating in the corresponding reaction. In the formula of the specific rates, we follow [3], noting that the formulas for the first three reactions is more complicated than a single-step enzyme reaction, since GA20ox participates in all of them.

$$\frac{\partial[\mathrm{GA}_j]}{\partial\hat{t}}+\frac{\partial}{\partial\hat{x}}\left(\hat{U}_{\mathrm{eff}}(\hat{x},\hat{t})[\mathrm{GA}_j]\right)=\frac{\partial}{\partial\hat{x}}\left(\hat{D}_{\mathrm{eff}}(\hat{x},\hat{t})\frac{\partial[\mathrm{GA}_j]}{\partial\hat{x}}\right)-\hat{Q}_{\mathrm{eff}}(\hat{x},\hat{t})[\mathrm{GA}_j]+\hat{S}_j \quad \text{for} \quad j=12,15,24,9,4, \tag{27}$$

where

$$\hat{S}_{12}=\frac{(1-\phi)\hat{w}\hat{l}\psi(\hat{x})}{\hat{V}}\left(\hat{S}(\hat{x})-\frac{\hat{\lambda}_{12}[\mathrm{GA}_{12}][\mathrm{GA20ox}]}{1+\hat{\kappa}_{12}[\mathrm{GA}_{12}]+\hat{\kappa}_{15}[\mathrm{GA}_{15}]+\hat{\kappa}_{24}[\mathrm{GA}_{24}]}-\hat{\beta}_{12}[\mathrm{GA}_{12}][\mathrm{GA2oxA}]\right), \tag{28a}$$

$$\hat{S}_{15}=\frac{(1-\phi)\hat{w}\hat{l}\psi(\hat{x})(\hat{\lambda}_{12}[\mathrm{GA}_{12}][\mathrm{GA20ox}]-\hat{\lambda}_{15}[\mathrm{GA}_{15}][\mathrm{GA20ox}])}{\hat{V}(1+\hat{\kappa}_{12}[\mathrm{GA}_{12}]+\hat{\kappa}_{15}[\mathrm{GA}_{15}]+\hat{\kappa}_{24}[\mathrm{GA}_{24}])}, \tag{28b}$$

$$\hat{S}_{24}=\frac{(1-\phi)\hat{w}\hat{l}\psi(\hat{x})(\hat{\lambda}_{15}[\mathrm{GA}_{15}][\mathrm{GA20ox}]-\hat{\lambda}_{24}[\mathrm{GA}_{24}][\mathrm{GA20ox}])}{\hat{V}(1+\hat{\kappa}_{12}[\mathrm{GA}_{12}]+\hat{\kappa}_{15}[\mathrm{GA}_{15}]+\hat{\kappa}_{24}[\mathrm{GA}_{24}])}, \tag{28c}$$

$$\hat{S}_{9}=\frac{(1-\phi)\hat{w}\hat{l}\psi(\hat{x})}{\hat{V}}\left(\frac{\hat{\lambda}_{24}[\mathrm{GA}_{24}][\mathrm{GA20ox}]}{1+\hat{\kappa}_{12}[\mathrm{GA}_{12}]+\hat{\kappa}_{15}[\mathrm{GA}_{15}]+\hat{\kappa}_{24}[\mathrm{GA}_{24}]}-\frac{\hat{\lambda}_{9}[\mathrm{GA}_{9}][\mathrm{GA3ox}]}{1+\hat{\kappa}_{9}[\mathrm{GA}_{9}]}-\hat{\beta}_{9}[\mathrm{GA}_{9}][\mathrm{GA2oxB}]\right), \tag{28d}$$

$$\hat{S}_{4}=\frac{(1-\phi)\hat{w}\hat{l}\psi(\hat{x})}{\hat{V}}\left(\frac{\hat{\lambda}_{9}[\mathrm{GA}_{9}][\mathrm{GA3ox}]}{1+\hat{\kappa}_{9}[\mathrm{GA}_{9}]}-\hat{\beta}_{4}[\mathrm{GA}_{4}][\mathrm{GA2oxC}]\right), \tag{28e}$$

[GA20ox], [GA3ox], [GA2oxA],[GA2oxB],[GA2oxC] are the corresponding enzyme transcript levels, which are given functions of $\hat{x}$ (see Table 3), where, based on current understanding of the GA2ox family [GA2oxB] = [GA2oxC], $\hat{S}(\hat{x})$ is the production rate of $GA_{12}$ (from either the QC or phloem pool, see (32) below), $\hat{D}_{\mathrm{eff}}, \hat{U}_{\mathrm{eff}}$, and $\hat{Q}_{\mathrm{eff}}$ are given in (8), $\hat{l}$, $\phi$, and $\hat{u}$ are given in (19), and the rate constants $\hat{\lambda}_j$ and $\hat{\kappa}_j$, $j = 12, 5, 24, 9, 4$, are given in Table 1.

In order to simplify (27) to compute numerical solutions, we expand the derivatives to obtain the simplified equations

$$\frac{\partial[\mathrm{GA}_j]}{\partial\hat{t}}=\hat{R}(\hat{x})(\hat{l}+\hat{a})\frac{\partial^2[\mathrm{GA}_j]}{\partial\hat{x}^2}+\left(\hat{R}(\hat{x})\frac{\partial\hat{l}}{\partial\hat{x}}-\hat{u}\right)\frac{\partial[\mathrm{GA}_j]}{\partial\hat{x}}-\frac{1}{\hat{V}}\frac{\mathrm{d}\hat{V}}{\mathrm{d}\hat{t}}[\mathrm{GA}_j]+\hat{S}_j \quad \text{for} \quad j=12,15,24,9,4, \tag{29}$$

where

$$\hat{R}(\hat{x})=\frac{(\hat{K}(\hat{l}+\hat{a})+\hat{M})}{\hat{V}}, \tag{30}$$

and $\hat{K}, \hat{M}$, and $\hat{V}$ are given in (9).

Equations (29) with (28a)–(28e) are solved separately in the meristem, elongation zone, and mature zone with (21) and (22), and are subject to the following boundary and initial conditions

$$\frac{\partial[\mathrm{GA}_j]}{\partial\hat{x}}=0 \quad \text{at} \quad \hat{x}=0, \tag{31a}$$

$$\frac{\partial[\mathrm{GA}_j]}{\partial\hat{x}}=0 \quad \text{at} \quad \hat{x}=\hat{L}_{\mathrm{root}}, \tag{31b}$$

$$[\mathrm{GA}_j]=0 \quad \text{at} \quad \hat{t}=0, \tag{31c}$$

where $j = 12, 15, 24, 9, 4$. Additionally, each $[\mathrm{GA}_j]$ and $\partial[\mathrm{GA}_j]/\partial\hat{x}$ are assumed to be continuous across the boundaries between the meristem, elongation zone, and the maturation zone, i.e., at $\hat{x} = \hat{L}_{\mathrm{meri}}$ and $\hat{x} = \hat{L}_{\mathrm{meri}} + \hat{L}_{\mathrm{ez}}$. We solve the time-dependent equations for large time when transport dynamics has stabilised.

We define the production rate of $GA_{12}$ to be

$$\hat{S}(\hat{x}) = \begin{cases} \hat{S}_{\mathrm{QC}} & \text{if } 0 \leq \hat{x} \leq \hat{L}_{\mathrm{QC}} \\ \hat{S}_{\mathrm{phloem}} & \text{if } \hat{L}_{\mathrm{meri}} \leq \hat{x} \leq \hat{L}_{\mathrm{phloem}} \\ 0 & \text{otherwise} \end{cases}, \tag{32}$$

where $\hat{L}_{\mathrm{phloem}}$ is the length of the phloem-unloading zone assumed to be half the length of the elongation zone, $\hat{S}_{\mathrm{QC}}$ is the rate of $GA_{12}$ production in the QCZ, and $\hat{S}_{\mathrm{phloem}}$ is the rate of $GA_{12}$ delivery in the phloem-unloading zone, respectively.

### 6 Parameter estimates

In Supplementary Tables 1–4, we list the model parameters together with the parameter values used in the simulations presented in the main text.

In Supplementary Tables 1-2, we list the physical model parameters with estimates of their typical values obtained from the experimental literature that pertain to the hormone gibberellin in the model species *Arabidopsis thaliana*. We note that, in order to obtain estimates for the transporter (NPF 3.1, NPF 2.14) permeabilities for the various metabolites, we used data, collected from experiments with oocytes placed in an external environment of a given metabolite concentration [24, 28]. These data are then fitted to a mathematical model, as in [6], to calculate the corresponding permeabilities. Due to difference in the structure of the transporters [28], the gibberellin metabolites can be grouped into fast ($GA_{12}$, $GA_{15}$, $GA_9$) and slowly ($GA_4$, $GA_{24}$) transported metabolites, and these two values appear in Tables 1-2.

Based on transcriptomics data [17, 20], we assume that the transcript levels of the enzymes are functions of the distance along the root, as shown in Fig. 1B of the main text. The values of the transcript levels of the enzymes, [GA20ox], [GA3ox], [GA2oxA], and [GA2oxB] are given in Supplementary Table 3. In order to obtain values for the rate constants of the metabolic reactions, we use the results in [27, 3].

Thus, the only unknown parameters in the model remain the synthesis/delivery rates of $GA_{12}$ and the degradation rates for $GA_4$, $GA_9$, and $GA_{12}$. In order to obtain estimates of their values, we performed a parameter survey, varying the synthesis rate in the QCZ, the delivery rate in the phloem-unloading zone, and the degradation rates separately and recording the square of the difference between the nlsGPS1 sensor data and the model prediction. We found that the best agreement between predictions and data corresponds to large degradation rates for $GA_{12}$, $GA_9$ and $GA_4$, and moderate synthesis/delivery rates (Supp. Fig. 4). This parameter survey led to the parameter values presented in Table 4.

### 7 Effect of cytoplasmic expansion in the elongation zone

We present a simple calculation, backed up by experimental data, to motivate our assumption that the cytoplasm continues to expand in the elongation zone.

The vacuolar fraction in the meristem is approximately $\phi_{\text{meri}} = 0.35$ [8]. Thus, to preserve this ratio, we must assume both the cytoplasm and the vacuole expand at the same rate, equal to the cell elongation rate in the meristem, $\hat{\text{RER}}_{\text{meri}}$. The cytoplasmic and total cellular volumes, $\hat{V}_{\text{cyt}}^{\text{meri}}$ and $\hat{V}_{\text{cell}}^{\text{meri}}$, respectively, in the meristem are therefore related by

$$\hat{V}_{\text{cyt}}^{\text{meri}} = (1 - \phi_{\text{meri}})\hat{V}_{\text{cell}}^{\text{meri}}. \tag{33}$$

In the elongation zone, cells increase their length (and therefore their volume) by approximately 10 times [9]. Thus, we assume

$$\hat{V}_{\text{cell}}^{\text{mat}} = 10\hat{V}_{\text{cell}}^{\text{meri}}, \tag{34}$$

where $\hat{V}_{\text{cell}}^{\text{mat}}$ is the cell volume at the end of the elongation zone (and the start of the maturation zone). If we further assume that, in the elongation zone, cell elongation is entirely due to vacuolar expansion, then the cytoplasmic volume remains the same, i.e., using (33) and (34), the cytoplasmic volume at the end of the elongation zone, $\hat{V}_{\text{cyt}}^{\text{mat}}$, is

$$\hat{V}_{\text{cyt}}^{\text{mat}} = \hat{V}_{\text{cyt}}^{\text{meri}} = (1 - \phi_{\text{meri}})\hat{V}_{\text{cell}}^{\text{meri}} = \frac{(1 - \phi_{\text{meri}})}{10}\hat{V}_{\text{cell}}^{\text{mat}} = 0.065, \tag{35}$$

which is smaller than the value, obtained experimentally in [7, 8].

If, now, we assume that the cytoplasm continues to expand at the same rate, as the cell enters the elongation zone, then

$$\frac{1}{\hat{V}_{\text{cyt}}^{\text{ez}}}\frac{\mathrm{d}\hat{V}_{\text{cyt}}^{\text{ez}}}{\mathrm{d}\hat{t}} = \hat{\text{RER}}_{\text{meri}}, \tag{36}$$

where $\hat{V}_{\text{cyt}}^{\text{ez}}$ is the cytoplasmic volume in the elongation zone, whereas the cell volume expands at the rate, equal to the elongation rate in the elongation zone, $\hat{\text{RER}}_{\text{ez}}$,

$$\frac{1}{\hat{V}_{\text{cell}}^{\text{ez}}}\frac{\mathrm{d}\hat{V}_{\text{cell}}^{\text{ez}}}{\mathrm{d}\hat{t}} = \hat{\text{RER}}_{\text{ez}}. \tag{37}$$

Dividing (36) by (37), we obtain

$$\frac{\mathrm{d}\hat{V}_{\text{cyt}}^{\text{ez}}}{\mathrm{d}\hat{V}_{\text{cell}}^{\text{ez}}} = \frac{\hat{\text{RER}}_{\text{meri}}}{\hat{\text{RER}}_{\text{ez}}}\frac{\hat{V}_{\text{cyt}}^{\text{ez}}}{\hat{V}_{\text{cell}}^{\text{ez}}}. \tag{38}$$

Integrating (38) and applying (33) and (34) gives

$$\hat{V}_{\text{cyt}}^{\text{mat}} = \frac{(1 - \phi_{\text{meri}})}{10^{1-\hat{\text{RER}}_{\text{meri}}/\hat{\text{RER}}_{\text{ez}}}}\hat{V}_{\text{cell}}^{\text{mat}} \approx 0.1. \tag{39}$$

This is precisely the value for the fraction of the cytoplasmic volume in the cell in the maturation zone in [7, 8].

### Supplementary figures

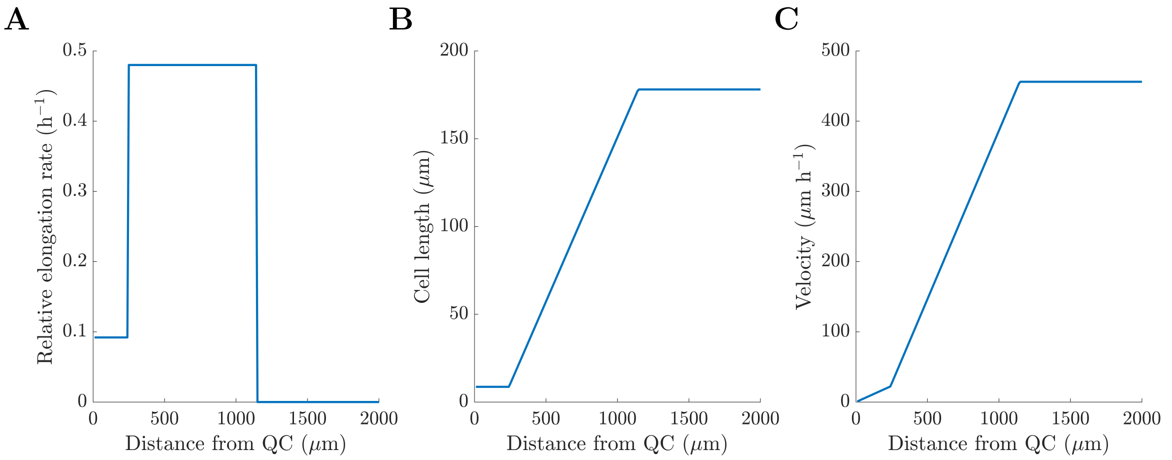


Supplementary Figure 1: **Prescribed cell RER, length, and velocity.** Spatial plots of the cell (**A**) relative elongation rate, (**B**) length, and (**C**) velocity.

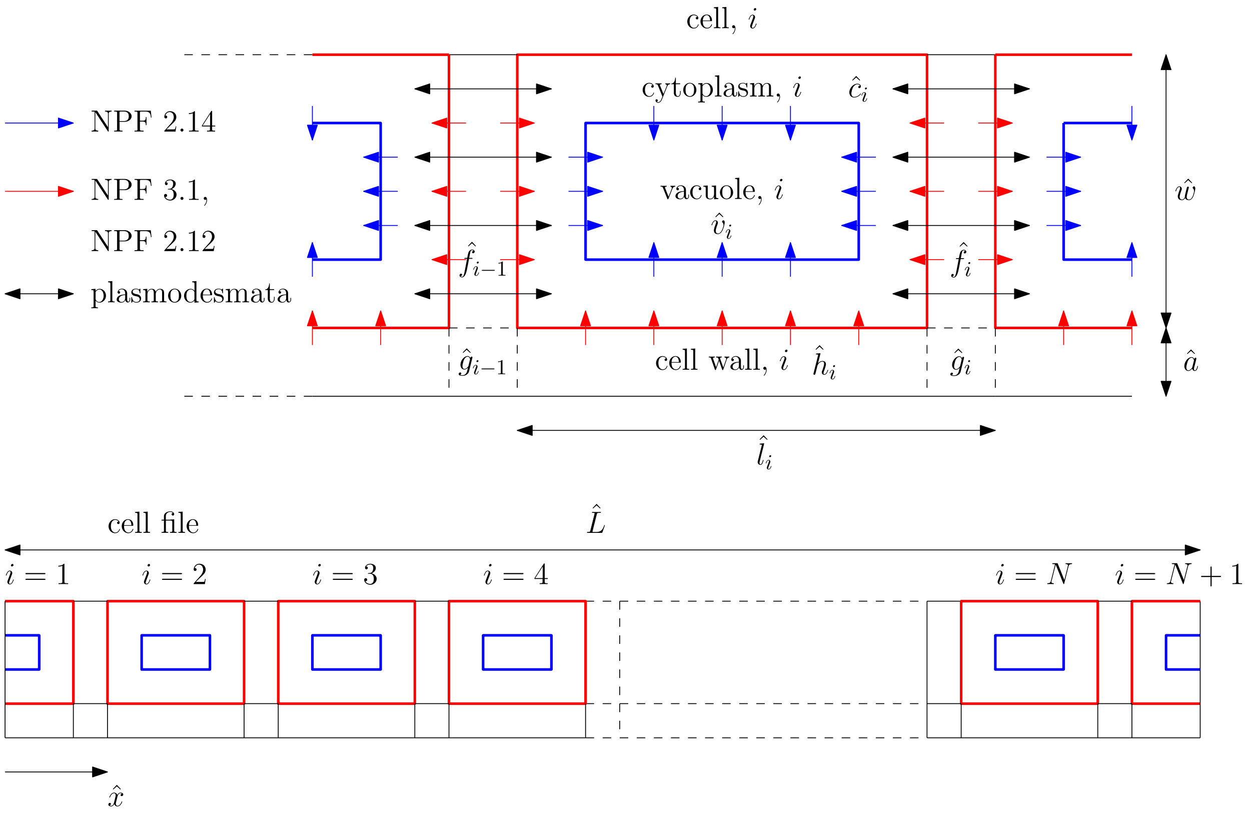


Supplementary Figure 2: Schematic of a file that consists of repeating cell units. Red arrows represent the cytoplasmic importers (NPF 2.12, 3.1 in the case of GA), whereas blue arrows represent the vacuolar importer (NPF 2.14 in the case of GA). The black arrows represent the plasmodesmata. The cytoplasmic membrane is shown in red, and the tonoplast is shown in blue [14].

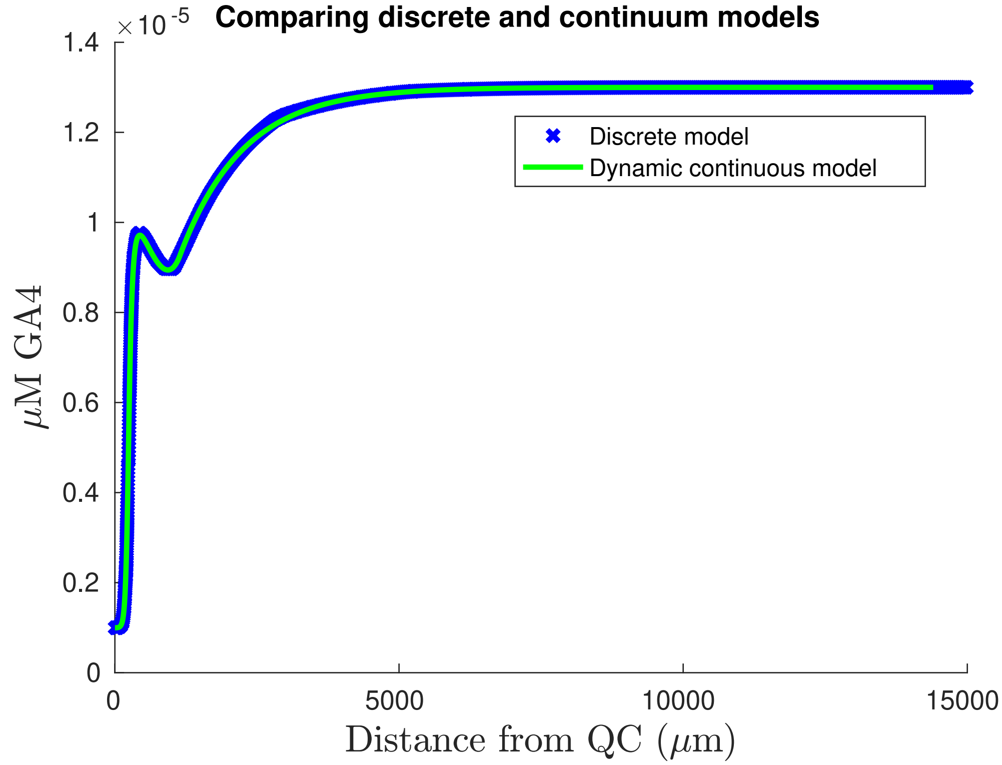


Supplementary Figure 3: **Continuum model agrees well with discrete model.** Comparison between the discrete cell-based and continuum model for $GA_4$ distribution, as detailed in section 3.

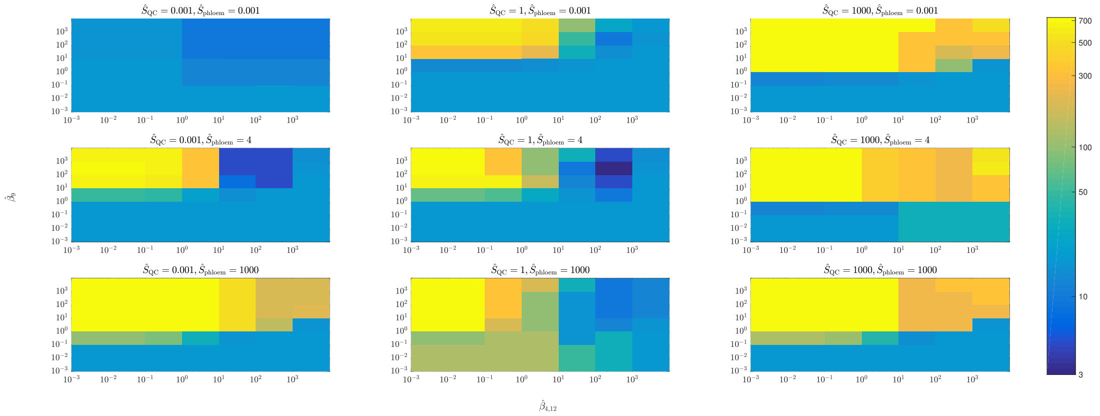

Supplementary Figure 4: **Sample figures from the outputs of the parameter survey showing the effect of the unknown parameter values on the agreement between model predictions and nlsGPS1 data.** The parameter survey considered 5-dimensional parameter space, surveying values of the five parameters: $\hat{S}_{QC}$, $\hat{S}_{phloem}$, $\hat{\beta}_{12}$, $\hat{\beta}_9$ and $\hat{\beta}_4$. Panels show slices through this 5-dimensional parameter space, with colours showing the square of the difference between the wildtype nlsGPS1 sensor data (shown in main text, Fig 2D) and the model prediction.

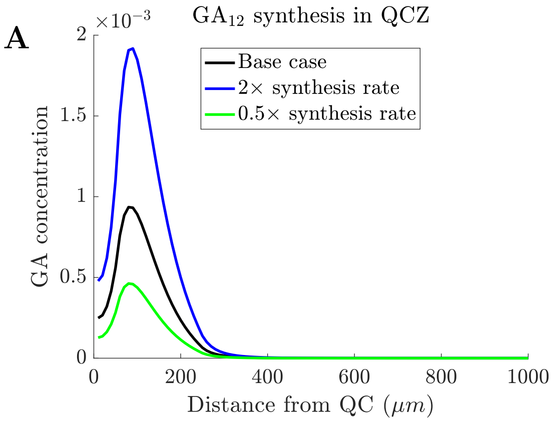


Supplementary Figure 5: **Varying synthesis/delivery rates of $GA_{12}$ scales the corresponding predicted $GA_4$ level.** (**A**) Predicted distribution of $GA_4$ originating from local $GA_{12}$ synthesis in the QCZ, with different values of the $GA_{12}$ synthesis rate. (**B**) Predicted distribution of $GA_4$ originating from phloem-delivered $GA_{12}$, with different values of the $GA_{12}$ delivery rate.

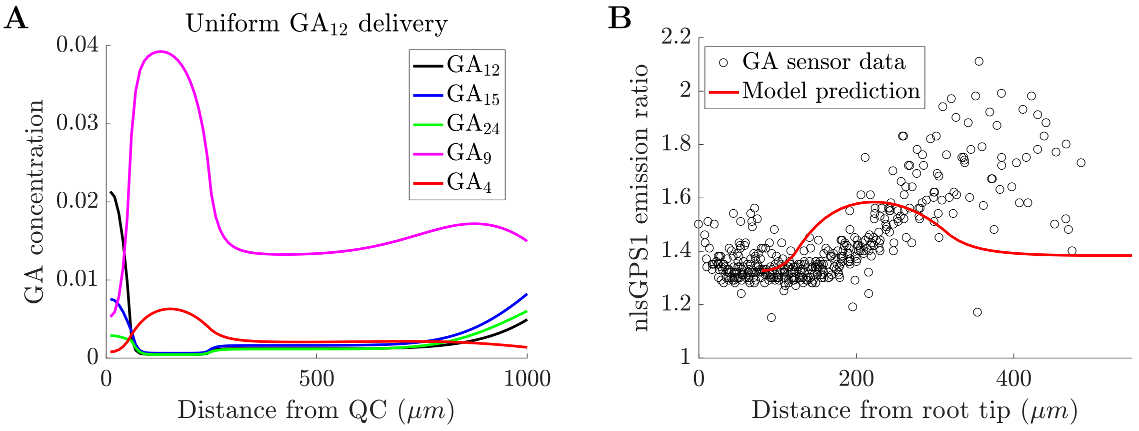


Supplementary Figure 6: **With uniform delivery/synthesis of $GA_{12}$, the model predictions do not reproduce the $GA_4$ gradient observed using the nls:GPS1 sensor.** (**A**) Predicted $GA_{12}$, $GA_{15}$, $GA_{24}$, $GA_9$, and $GA_4$ distributions with uniform $GA_{12}$ delivery. (**B**) Comparison between model predictions and experimental data for the nlsGPS1 emission ratio.

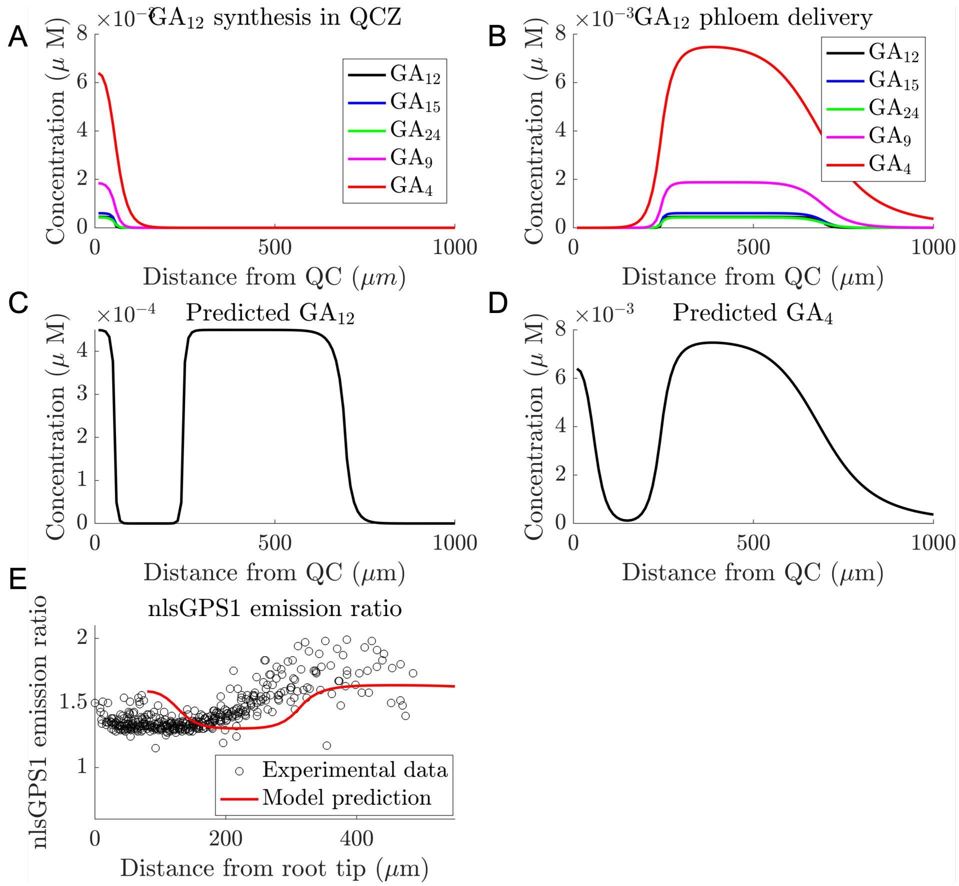


Supplementary Figure 7: **With uniform enzyme transcript levels, the model predictions do not reproduce the $GA_4$ gradient observed using the nls:GPS1 sensor.** (**A,B**) Predicted $GA_{12}$, $GA_{15}$, $GA_{24}$, $GA_9$, and $GA_4$ distributions with uniform enzyme levels due to (**A**) $GA_{12}$ synthesis in the QCZ and (**B**) $GA_{12}$ delivery the phloem-unloading zone. (**C**) Predicted $GA_{12}$ distribution (due to both $GA_{12}$ delivery and local synthesis). (**D**) Predicted $GA_4$ distribution (due to both $GA_{12}$ delivery and local synthesis). (**E**) Comparison between model predictions and experimental data for the nlsGPS1 emission ratio. Predicted sensor distribution is calculated from the base case $GA_4$ distribution shown in panel D. Parameter values are given in Tables 1 and 4, and the enzyme levels are set to unity: [GA20ox] = [GA3ox] = [GA2oxA] = [GA2oxB] = 1.

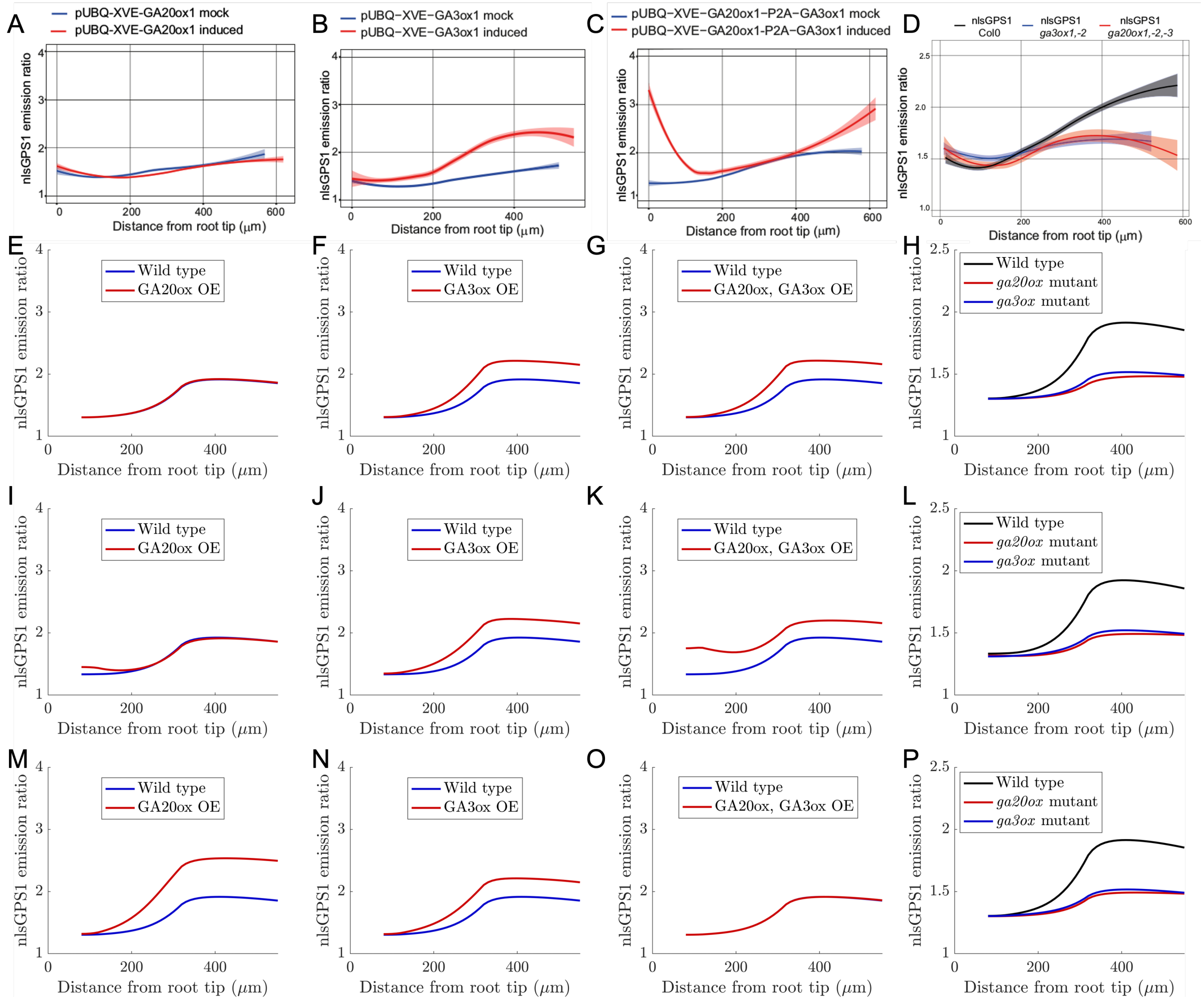


Supplementary Figure 8: **Alternative assumptions about enzyme actvity do not lead to model predictions that are consistent with the nlsGPS1 data in the GA20ox and GA3ox over-expression lines.** (**A-D**) Experimental nlsGPS1 data reproduced from [20]. (**E-P**) Predicted nls-GPS1 emission ratio for (**E,I,P**) GA20ox overexpression, (**F,J,N**) GA3ox overexpression, (**G,K,O**) both GA20ox and GA3ox overexpression, and (**H,L,P**) *ga20ox* and *ga3ox* loss-of-function mutants. (**E-H**) Model predictions in which the GA20ox and GA3ox activity is set to zero throughout the meristem. (**I-L**) Model predictions in which the GA20ox and GA3ox activity is reduced by 10 times in the DZ. (**I-L**) Model predictions in which the GA20ox and GA3ox activity is reduced by 10 times throughout the meristem. In each case, all other enzyme activities are based on the transcript level (shown in main text Fig 1B).

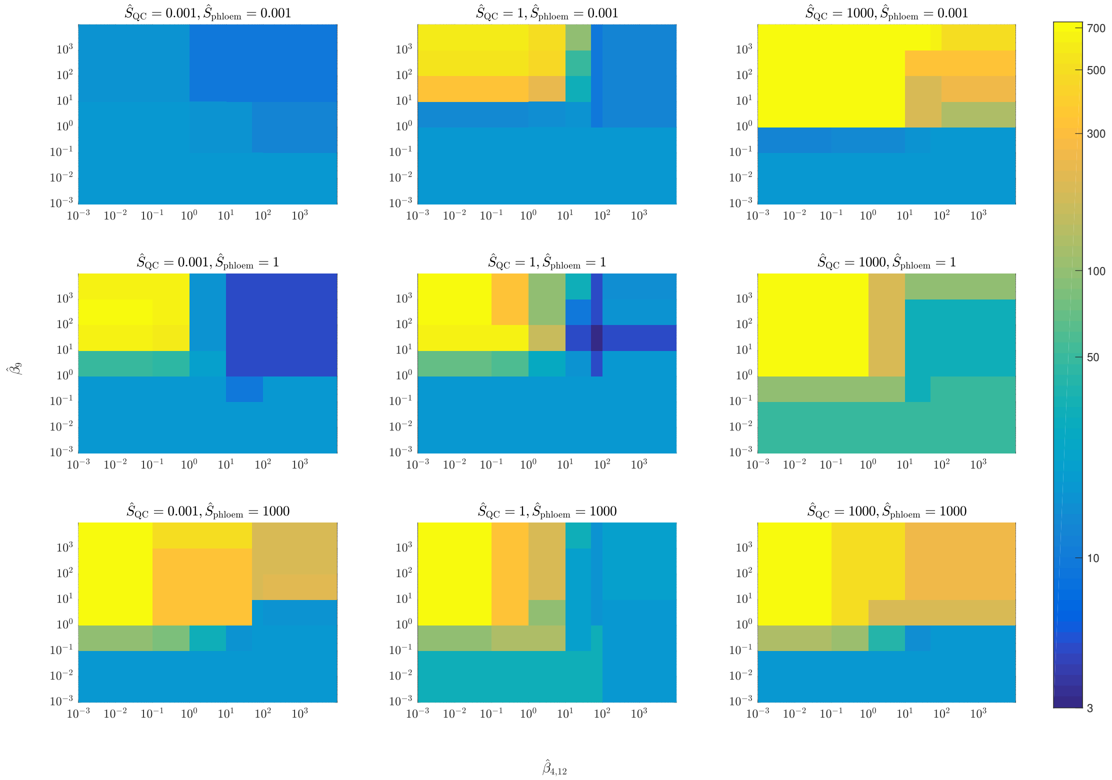


Supplementary Figure 9: **Sample figures from the outputs of the parameter survey showing the effect of the unknown parameter values on the agreement between model predictions and nlsGPS1 data, assuming that GA20ox and GA3ox are inactive in the DZ.** The parameter survey considered 5-dimensional parameter space, surveying values of the five parameters: $\hat{S}_{QC}$, $\hat{S}_{phloem}$, $\hat{\beta}_{12}$, $\hat{\beta}_9$ and $\hat{\beta}_4$, in the range $10^{-3}$ and $10^3$. Figures show nine slices through this 5-dimensional parameter space. Colours show the square of the difference between the wildtype nlsGPS1 sensor data (shown in main text, Fig 2D) and the model prediction for each parameter set shown.

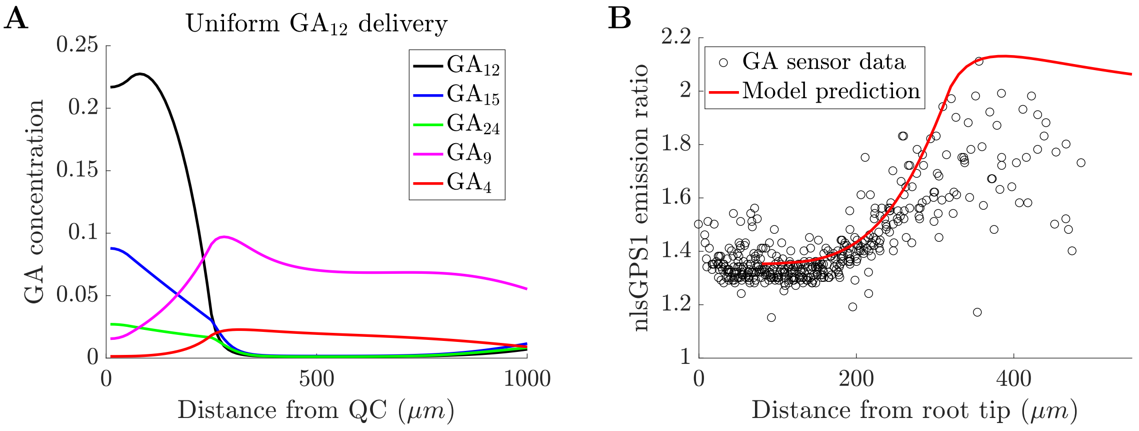


Supplementary Figure 10: **With uniform delivery/synthesis of $GA_{12}$ and GA20ox and GA3ox enzyme activities set to zero in the DZ, the model predictions are able to approximate the $GA_4$ gradient observed using the nls:GPS1 sensor.** (**A**) Predicted $GA_{12}$, $GA_{15}$, $GA_{24}$, $GA_9$, and $GA_4$ distributions with uniform $GA_{12}$ synthesis/delivery. (**B**) Comparison between model predictions and experimental data for the nlsGPS1 sensor emission ratio.

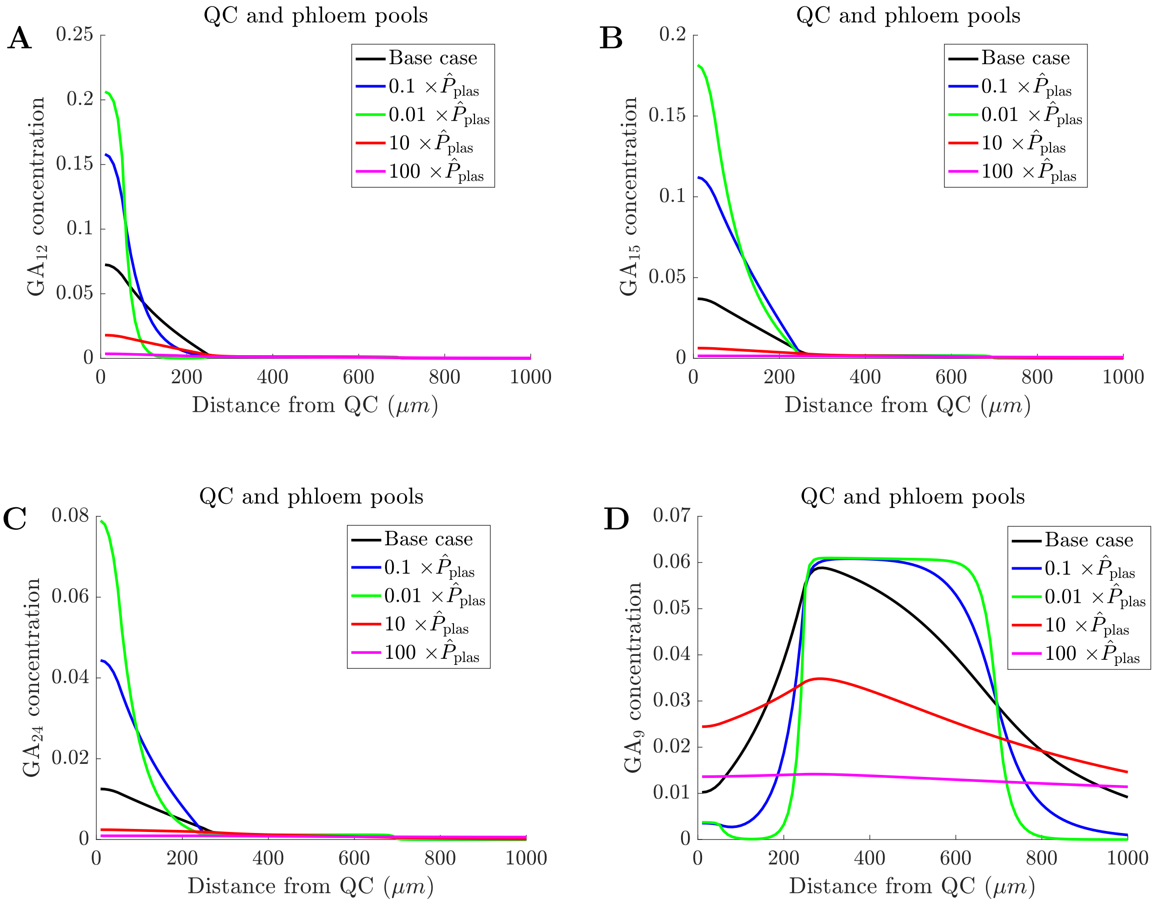


Supplementary Figure 11: **Plasmodesmata have a significant effect on GA gradient.** Predicted (**A**) $GA_{12}$, (**B**) $GA_{15}$, (**C**) $GA_{24}$, and (**D**) $GA_9$ distributions with varying plasmodesmata permeability.

### Supplementary tables

| Parameter | Description | Value | Reference |
|---|---|---|---|
| $\hat{l}_0$ | initial cell length in meristem | $6\,\mu$m | [9] |
| $\hat{L}_{\text{root}}$ | file length | $1300\,\mu$m | [20] |
| $\hat{L}_{\text{QC}}$ | length of quiescent centre zone | $43.5\,\mu$m | [18] |
| $\hat{L}_{\text{phloem}}$ | length of phloem-unloading zone | $450\,\mu$m | [21] |
| $\hat{L}_{\text{meri}}$ | length of meristem | $240\,\mu$m | [20] |
| $\hat{L}_{\text{ez}}$ | length of elongation zone | $900\,\mu$m | [20] |
| $\phi_{\text{meri}}$ | vacuolar fraction in meristem | 0.35 | [8] |
| $\phi_{\text{mat}}$ | vacuolar fraction in maturation zone | 0.9 | [8] |
| $\hat{\text{RER}}_{\text{meri}}$ | cell elongation rate in meristem | $0.092\,\text{h}^{-1}$ | [25] |
| $\hat{\text{RER}}_{\text{ez}}$ | cell elongation rate in elongation zone | $0.48\,\text{h}^{-1}$ | [25] |
| $\hat{w}$ | cell width | $10\,\mu$m | [2] |
| $\hat{a}$ | apoplast thickness | $0.5\,\mu$m | [2] |
| $\hat{D}_{\text{apo}}$ | apoplastic diffusivity | $32\,\mu\text{m}^2\text{s}^{-1}$ | [16] |
| $\hat{P}_{\text{plas}}$ | plasmodesmatal permeability | $0.81\,\mu\text{m}\,\text{s}^{-1}$ | [22] |
| $\hat{\lambda}_{12}$ | rate of conversion from $GA_{12}$ to $GA_{15}$ | $2580\,\mu\text{M}^{-1}\text{h}^{-1}$ | [3] |
| $\hat{\lambda}_{15}$ | rate of conversion from $GA_{15}$ to $GA_{24}$ | $1920\,\mu\text{M}^{-1}\text{h}^{-1}$ | [3] |
| $\hat{\lambda}_{24}$ | rate of conversion from $GA_{24}$ to $GA_9$ | $2700\,\mu\text{M}^{-1}\text{h}^{-1}$ | [3] |
| $\hat{\lambda}_9$ | rate of conversion from $GA_9$ to $GA_4$ | $408\,\mu\text{M}^{-1}\text{h}^{-1}$ | [27] |
| $\hat{\kappa}_{12}$ | rate of conversion from $GA_{12}$ to $GA_{15}$ | $0\,\mu\text{M}^{-1}\text{h}^{-1}$ | [3] |
| $\hat{\kappa}_{15}$ | rate of conversion from $GA_{15}$ to $GA_{24}$ | $0\,\mu\text{M}^{-1}\text{h}^{-1}$ | [3] |
| $\hat{\kappa}_{24}$ | rate of conversion from $GA_{24}$ to $GA_9$ | $500\,\mu\text{M}^{-1}\text{h}^{-1}$ | [3] |
| $\hat{\kappa}_9$ | rate of conversion from $GA_9$ to $GA_4$ | $1\,\mu\text{M}^{-1}\text{h}^{-1}$ | [27] |
| $\hat{P}_{\text{ca}}$ | $B_1\hat{P}_{\text{pass}} + B_2\hat{P}_{\text{imp}}$ | $0.0064/0.0054\,\mu\text{m}\,\text{s}^{-1}$ | [$\star$] |
| $\hat{P}_{\text{ac}}$ | $A_1\hat{P}_{\text{pass}} + A_2\hat{P}_{\text{imp}}$ | $0.629/0.714\,\mu\text{m}\,\text{s}^{-1}$ | [$\star$] |
| $\hat{P}_{\text{cv}}$ | $B_1\hat{P}_{\text{pass}} + B_3\hat{P}_{\text{exp}}$ | $0.940/0.0493\,\mu\text{m}\,\text{s}^{-1}$ | [$\star$] |
| $\hat{P}_{\text{vc}}$ | $C_1\hat{P}_{\text{pass}} + C_3\hat{P}_{\text{exp}}$ | $0.295/0.159\,\mu\text{m}\,\text{s}^{-1}$ | [$\star$] |

Supplementary Table 1: Physical model parameter estimates obtained from the cited literature. The quantities with stars [$\star$] are calculated from the quantities and formulae listed in the Table 2. Left-hand-side estimates pertain to $GA_4$ and $GA_{24}$, whereas the corresponding estimates on the right-hand side pertain to $GA_{12}$, $GA_{15}$, and $GA_9$.

| Parameter | Description | Value | Reference |
|---|---|---|---|
| $pH_{cyt}$ | cytoplasmic pH | 7.6 | [23] |
| $pH_{apo}$ | apoplastic pH | 5.3 | [1] |
| $pH_{vac}$ | vacuolar pH | 5.8 | [12] |
| pK | equilibrium dissociation constant | 4.2/4.3 | [15] |
| $\hat{D}_{apo}$ | apoplastic diffusivity | $32\,\mu m^2 s^{-1}$ | [16] |
| $V_{mem}$ | potential across cell membrane | $-120\,mV$ | [1, 23] |
| $V_{ton}$ | potential across tonoplast | $-30\,mV$ | [11, 23] |
| $T$ | absolute temperature | $300\,K$ | [1] |
| $\hat{P}_{pass}$ | passive permeability | $0.333/4.72\,\mu m\,s^{-1}$ | [24, 6] |
| $\hat{P}_{imp}$ | importer permeability | $0.139/0.0667\,\mu m\,s^{-1}$ | [24, 6] |
| $\hat{P}_{exp}$ | exporter permeability | $0.556/0.0278\,\mu m\,s^{-1}$ | [28, 6] |
| $F_D$ | Faraday's constant | $96500\,C\,mol^{-1}$ | [1] |
| $R$ | universal gas constant | $8.31\,J\,mol^{-1}K^{-1}$ | [1] |
| $A_1$ | $1/(1+10^{pH_{apo}-pK})$ | 0.0736/0.0909 | [$\star$] |
| $A_2$ | $(F_D\hat{V}_{mem}/R\hat{T})(1-A_1)/(e^{F_D\hat{V}_{mem}/R\hat{T}}-1)$ | 4.34/4.26 | [$\star$] |
| $B_1$ | $1/(1+10^{pH_{cyt}-pK})$ | 0.000501/0.000398 | [$\star$] |
| $B_2$ | $-(F_D\hat{V}_{mem}/R\hat{T})(1-B_1)/(e^{-F_D\hat{V}_{mem}/R\hat{T}}-1)$ | 0.0451/0.045 | [$\star$] |
| $B_3$ | $(F_D\hat{V}_{ton}/R\hat{T})(1-B_1)/(e^{F_D\hat{V}_{ton}/R\hat{T}}-1)$ | 1.69/1.69 | [$\star$] |
| $C_1$ | $1/(1+10^{pH_{vac}-pK})$ | 0.0245/0.0307 | [$\star$] |
| $C_3$ | $-(F_D\hat{V}_{ton}/R\hat{T})(1-C_1)/(e^{-F_D\hat{V}_{ton}/R\hat{T}}-1)$ | 0.516/0.513 | [$\star$] |

Supplementary Table 2: Physical parameter estimates used in calculating the model parameter values related to transport (listed and marked with stars in Table 1). The quantities with stars here [$\star$] are calculated from other quantities in this table according to the corresponding formula. The permeability estimates are obtained from oocyte experiments, as in our previous work [6]. Their values, the pK values, and the related constants on the left-hand side pertain to $GA_4$ and $GA_{24}$, whereas the corresponding estimates on the right-hand side pertain to $GA_{12}$, $GA_{15}$, and $GA_9$. Here, for concreteness, we have taken $\hat{P}_{imp}$ to refer to the permeability estimated from oocyte experiments with NPF3.1. $\hat{P}_{exp}$ refers to the permeability estimated from data for NPF2.14.

| Region | [GA20ox] | [GA3ox] | [GA2oxA] | [GA2oxB, C] |
|---|---|---|---|---|
| quiescent-centre zone (QCZ) | 0.01 | 0.01 | 0.08 | 1 |
| division zone (DZ) | 1 | 0.01 | 0.04 | 0.2 |
| elongation zone (EZ) | 0.5 | 0.025 | 0.08 | 0.65 |
| maturation zone (MZ) | 0.05 | 0.0025 | 0.08 | 10 |

| Region | [GA20ox] | [GA3ox] | [GA2oxA] | [GA2oxB] |
|---|---|---|---|---|

Supplementary Table 3: Enzyme transcript levels assumed in the model, based on transcriptomics data in [20, 17]. [GA20ox] and [GA3ox] in the DZ are set to zero or reduced to 0.1× in simulations with reduced biosynthesis enzyme activity in the DZ.

| Parameter | Description | Value |
|---|---|---|
| $\hat{S}_{\mathrm{QC}}$ | production rate of $GA_{12}$ in QCZ | $1\,\mu\mathrm{M}\,\mathrm{h}^{-1}$ |
| $\hat{S}_{\mathrm{phloem}}$ | delivery rate of $GA_{12}$ in phloem-unloading zone | $4/1\,\mu\mathrm{M}\,\mathrm{h}^{-1}$ |
| $\hat{\beta}_{12}$ | rate of degradation of $GA_{12}$ | $100/50\,\mathrm{h}^{-1}$ |
| $\hat{\beta}_{9}$ | rate of degradation of $GA_9$ | $100/10\,\mathrm{h}^{-1}$ |
| $\hat{\beta}_{4}$ | rate of degradation of $GA_4$ | $100/50\,\mathrm{h}^{-1}$ |

Supplementary Table 4: Parameter estimates obtained from the parameter survey. The left-hand values are for the initial simulations where enzyme activities depend on the transcript level, and the right-hand values are for the simulations with reduced activity of GA20ox and GA3ox in the DZ.

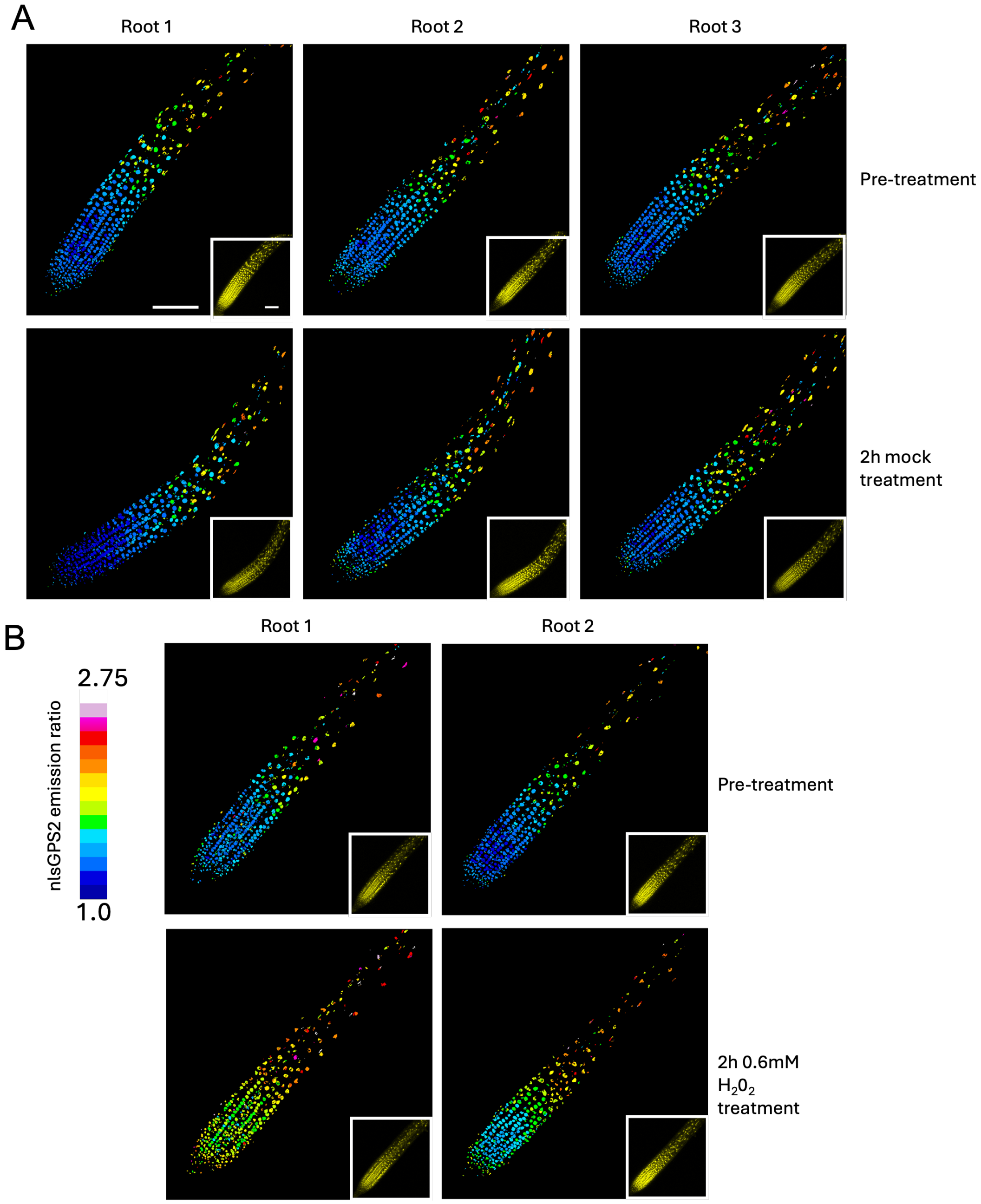


Supplementary Figure 12: **Additional images from $H_20_2$ treatment experiments.** Images of Col-0 nlsGPS2 roots. Representative images of emission ratios and YFP fluorescence (Inset) are shown of A) pre-treated and 2h mock-treated roots. B) pre-treated and 2h 0.6mM $H_20_2$ treated roots. Colour bar and scale bar apply to all images. Scale bar = 100$\mu$m.

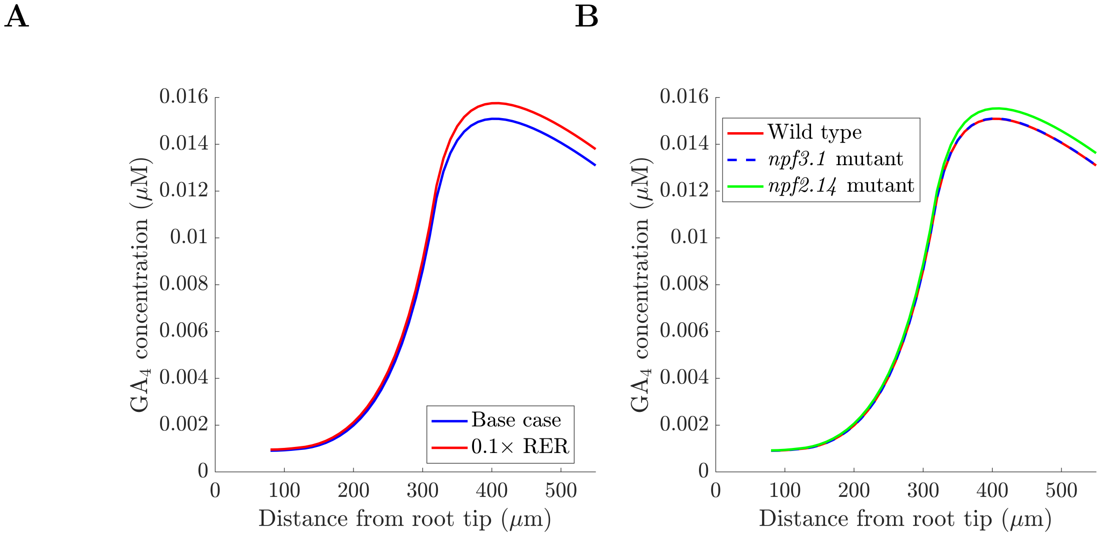


Supplementary Figure 13: **Dilution and NPF-mediated transport have limited effect on the predicted $GA_4$ distribution.** (**A**) Predicted $GA_4$ distribution with the relative elongation rates reduced by ten times (i.e. $\hat{\mathrm{RER}}_{\mathrm{meri}} = 0.0092$ and $\hat{\mathrm{RER}}_{\mathrm{EZ}} = 0.048$). (**B**) Predicted $GA_4$ distribution in wildtype, the *npf3.1* mutant ($\hat{P}_{\mathrm{imp}} = 0$) and the *npf2.14* mutant ($\hat{P}_{\mathrm{exp}} = 0$). In each case, the remaining parameter values are given in Tables 1, 2 and 4.

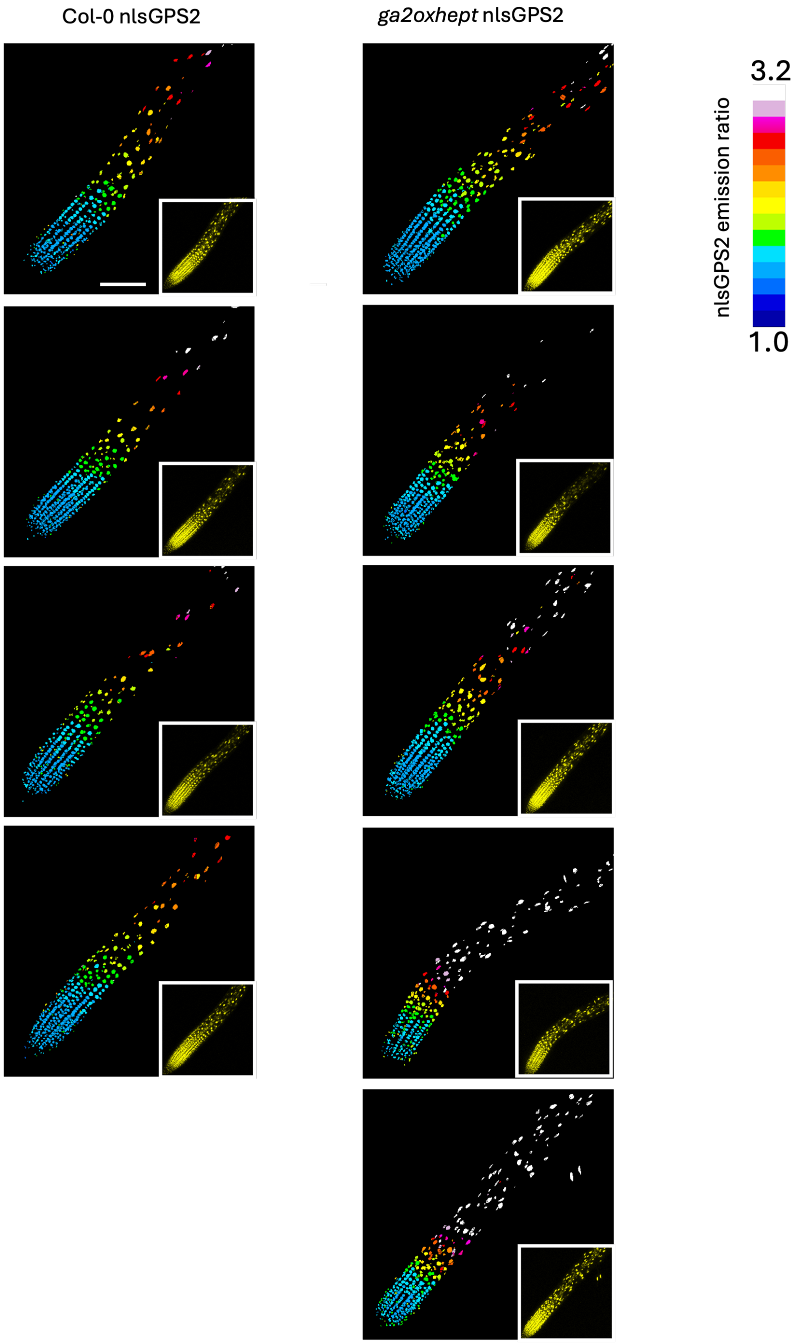


Supplementary Figure 14: **Additional images of Col-0 nlsGPS2 and ga2oxhept nlsGPS2 roots.** Representative images of emission ratios and YFP fluorescence (Inset) are shown. Colour bar and scale bar apply to all images. Scale bar = 100$\mu$m.

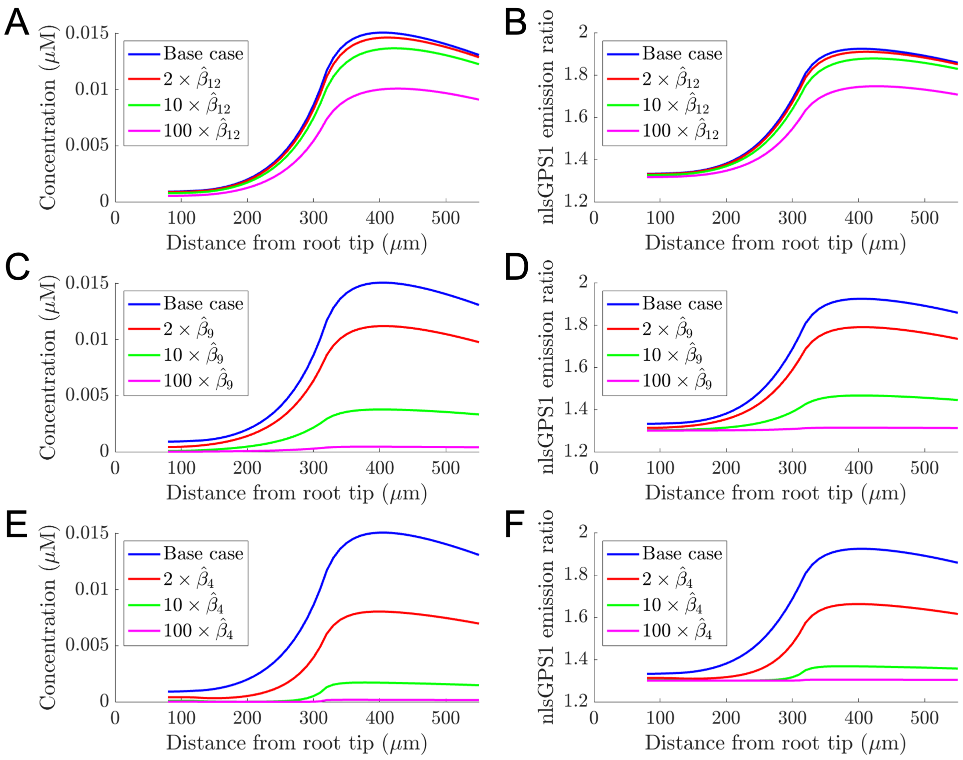


Supplementary Figure 15: **Effect of increasing the GA2ox-mediated degradation rates on the predicted $GA_4$ and nlsGPS1 distributions.** (**A, B**) Model predictions increasing the $GA_{12}$ degradation rate, $\hat{\beta}_{12}$. (**C,D**) Model predictions increasing the $GA_9$ degradation rate, $\hat{\beta}_9$. (**E,F**) Model predictions increasing the $GA_4$ degradation rate, $\hat{\beta}_4$. We note that increasing a degradation rate to 10× or 100× would represent the corresponding GA2ox overexpression line.
